# GSET: Gene Signals from Embedding Trajectories select compact contexts for mechanistic GRN inference

**DOI:** 10.64898/2026.10.30.763241

**Authors:** Zeev Weizmann, Elias Ventre

## Abstract

Mechanistic inference of gene regulatory networks from temporal transcriptomic data can become computationally prohibitive at transcriptome scale, making gene selection an important determinant of which regulatory relationships can be recovered. However, gene selection for GRN inference commonly relies on static expression or co-expression structure, rather than prioritizing dynamically informative genes whose temporal patterns capture how transcriptional programs change and relate to one another. We introduce GSET (Gene Signals from Embedding Trajectories), an upstream retrieval framework that extracts dynamic information from temporal gene embeddings using two complementary signals: an optimal transport residual, which captures gene-specific deviations from global embedding movement, and a KL divergence, which detects changes in local co-expression neighborhoods. Temporal gene-embedding similarity identifies static transcriptional programs, whereas vectorial GSET signals represent coordinated temporal changes among genes, complementing this static structure. Lagged comparisons of these signals further prioritize temporal relationships between programs and candidate bridge genes associated with their transitions. Integrated with CardamomOT, GSET combines static program cores with dynamically retrieved bridge genes into compact, program-informed gene sets for mechanistic GRN inference. In synthetic benchmarks, at equal gene budget these sets recover more of the regulatory links between programs than static selections of the same size, and inference on the edges within the set is comparable to that of the full-gene model on the same gene pairs. On a time course of mouse fibroblasts reprogrammed into induced pluripotent stem cells, GSET traces the transition from the fibroblast program to the cell-cycle program. By providing a dynamic retrieval layer upstream of GRN inference, GSET nominates bridge genes as candidate regulators of cell-state transitions, which, once refined by a mechanistic model such as CardamomOT and confirmed by perturbation experiments, could open the way to controlling these transitions.

## 1 Introduction

Mechanistic and dynamical models of gene regulatory networks (GRNs) fitted to time-resolved single-cell data are emerging as a promising way to recover the direction and sign of regulatory interactions [13, 37, 38, 43, 2, 29, 35, 23]. However, the number of candidate interactions grows quadratically with the number of genes, which increases both the computational cost of these models and the number of parameters to be estimated from a fixed number of cells. Which genes enter the model therefore largely determines which regulatory relationships can be recovered. Current gene selection is largely static: genes are ranked by their variability, by membership in curated transcription-factor lists [26], or by static co-expression [19]. The selected genes distinguish cell states [28], but reveal neither how a state is actively maintained nor how cells exit it towards the next.

A cell state is held in place by coordinated gene activity [39, 10]. Genes act in concert to maintain a cell’s identity, to respond to external signals and to carry out activities such as replication and metabolism [16]. This coordination often arises from co-regulation, with genes induced together by a shared internal or external signal [6, 32], a notion that Kotliar et al. [16] formalised operationally as a gene expression program. Maintaining a cell state or exiting it therefore corresponds to sustaining or switching the programs that underlie it.

Programs can be represented through the structural signature that co-regulation leaves in co-expression. This signature can be captured by a co-expression graph whose nodes are genes and whose edges encode pairwise co-expression strength, for instance the soft-thresholded correlation weights of WGCNA [19]. Graph neural networks, such as graph autoencoders [14], can then encode this graph into gene embeddings, an approach already used for gene-module and gene-interaction analysis [41, 20]. Because such encoders place strongly co-expressed genes close together, genes of the same program form compact neighbourhoods in the latent space.

Such an embedding describes programs at a single timepoint but does not show how they change. A program can change in two ways: its genes can shift together in activity while keeping the same co-expression partners, or a gene can acquire new partners without any change in its own expression level. With time-resolved data, genes can be embedded at every timepoint, so that each gene traces a trajectory in latent space that can reveal both kinds of change. Not every movement is informative, however: the whole embedding can drift between timepoints because of population-level responses or technical effects. Separating a gene’s own movement from this global drift is analogous to decomposing an asset’s return into a market component and an idiosyncratic component [33].

A second difficulty concerns how changes in different programs relate to one another. Regulation that *maintains* a program and regulation that drives *exit* from it are expected to appear as co-change and as temporal precedence, respectively, but which signature a given relationship produces depends on the sampling interval relative to its biological delay, and sampling grids are rarely matched to these timescales.

Because the selected genes are only an input to a downstream mechanistic model, the requirements on selection differ from those of a stand-alone analysis. First, it must scale to the whole transcriptome, since its purpose is to avoid running the mechanistic model at that scale. Second, programs must be identified *consistently*, across seeds and queries. Which of two genes with near-identical dynamics represents a program matters little, since either carries the same information. Third, the costs of errors are asymmetric: a spurious bridge gene only enlarges the gene budget of the downstream model, whereas a missed bridge removes a regulatory link between programs that no downstream inference can recover. Selection should therefore favour recall of bridges over precision.

GSET (Gene Signals from Embedding Trajectories) addresses these difficulties in turn. It follows each gene along two embedding trajectories. A change of co-expression partners is visible in the co-expression embedding. Because this change is defined by the relative neighbourhood structure rather than by absolute latent coordinates, GSET measures it with a symmetrised KL divergence [18] between a gene’s neighbourhood distributions at successive timepoints. A change in activity is not directly visible there, because message passing and the scale invariance of correlation discard each gene’s own expression level. GSET therefore embeds each gene’s marginal expression distribution separately, estimates the global drift with optimal transport (OT) [3, 31], and keeps each gene’s residual. The vectorial forms of both signals reveal coordinated gene-level temporal change, complementing the static program structure recovered from the underlying embeddings. To cope with unknown timescales, GSET then uses channels at different temporal lags, from co-change to precedence, with a fixed pair of channels chosen from a comparison of lags across sampling grids rather than tuned to a dataset. Around any query gene, the resulting relationships retrieve a compact, dynamically informative gene set, which is exactly what the mechanistic models above require. GSET thus acts as a retrieval layer [21] and leaves the inference of causal edges to the downstream model.

### Related work

Gene program methods such as cNMF [16], Hotspot [5] and PRISM-GEP [1], which allows overlapping programs, identify programs from static co-expression. GeneTrajectory [27] orders genes along a trajectory using the OT-based gene–gene distance of Huizing et al. [11] over a static cell–cell kNN graph that pools all timepoints, OTVelo [43] infers lagged gene–gene relations from OT couplings between the *cells* of consecutive snapshots, and DELVE [28] selects features that co-vary smoothly along a cell-similarity graph, without using sampling time. None of these recovers ordered temporal relationships between programs: cNMF and Hotspot omit time, PRISM-GEP orders genes along a diffusion coordinate rather than by measured time, GeneTrajectory and OTVelo operate on genes and gene pairs rather than programs, and DELVE substitutes trajectory similarity for measured time. GSET instead works on the measured timepoints and transports *genes* rather than cells: after embedding, every OT and KL computation takes place in gene space, with one embedded point per gene and timepoint, so it needs no cell–cell couplings or gene–gene OT problems over cells. Time-series GRN methods such as RENGE [13], FlowGRN [35] and CardamomOT [23] infer edges directly but are typically run on a pre-selected gene panel, and GSET is a selection layer that decides which genes such a method should see.

### Contributions

- Two gene-level signals of temporal change: an OT residual in a marginal-expression embedding, and a KL neighbourhood divergence in a co-expression embedding. In controlled simulations they respond preferentially to activity shifts and to co-expression rewiring, respectively, and, in the scenario that suits each signal, they preserve program structure that naive differencing loses.
- A retrieval layer that combines program cores with bridge genes selected by lagged comparison of these signals. KL at lag 0 carries most of the retrieval, and OT at lag 1 gives direction information beyond program order. The best OT lag shifts with sampling resolution, which motivates fixed channels rather than tuning a lag to each dataset.
- An integration with CardamomOT. At equal gene budget, GSET contexts cover more relay edges between programs than static selections, with no loss on the edges inside the context compared with the full-gene model, at a cost that keeps mechanistic inference feasible on thousands of genes. Real-data checks on reprogramming and rhabdomyosarcoma time courses complement the synthetic benchmarks.

## 2 Methods

### 2.1 Overview

The data are single-cell snapshots of *n* genes at timepoints *t*_0_ < · · · < *t*_*M*_. Sequencing destroys the cells, so each timepoint measures different cells and no cell can be followed over time. From these snapshots, GSET (Fig. 1) (i) embeds every gene at every timepoint in a co-expression space and a marginal-expression space (§2.2), (ii) computes the per-transition signals *δ*^KL^ and *δ*^OT^ (§2.3–2.4), (iii) identifies static programs from concatenated co-expression embeddings (§2.5), (iv) compares the signal trajectories across lags to score co-change and statistical precedence and to retrieve bridge genes between programs (§2.6), and (v) combines program cores with bridges into a compact context for CardamomOT (§2.7).

**Figure 1:**
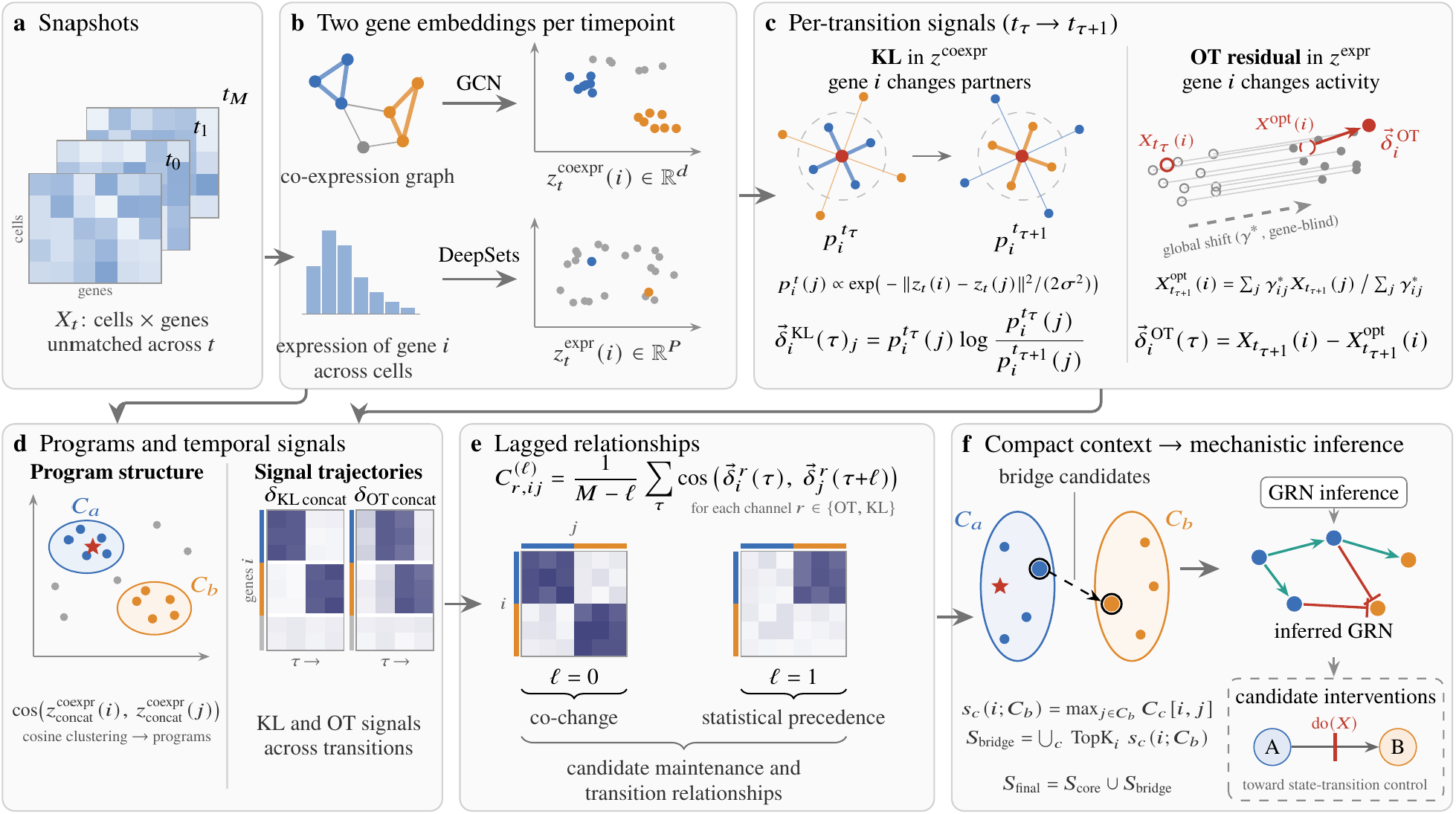
GSET overview. **(a)** Time-resolved snapshots, with cells unmatched across timepoints. **(b)** At each timepoint, a shared-weight GCN embeds genes by co-expression 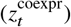 and a DeepSets encoder by their expression distribution 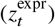. **(c)** For each transition, 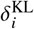 measures the change of gene *i*’s co-expression neighbourhood and 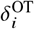 its displacement after removing the global OT shift. **(d)** Concatenated co-expression embeddings define static programs (*C*_*a*_, *C*_*b*_; star: query gene), and the signals form gene-level trajectories across transitions (rows: genes, coloured by program; columns: transitions). **(e)** Lagged cosine comparisons score co-change (ℓ= 0) and precedence (ℓ≥ 1). **(f)** Bridge genes retrieved across channels are combined with program cores into the context *S*_core_ ∪ *S*_bridge_, on which CardamomOT infers a directed, signed network. The dashed box indicates a prospective use that is not evaluated here.

**Figure 2:**
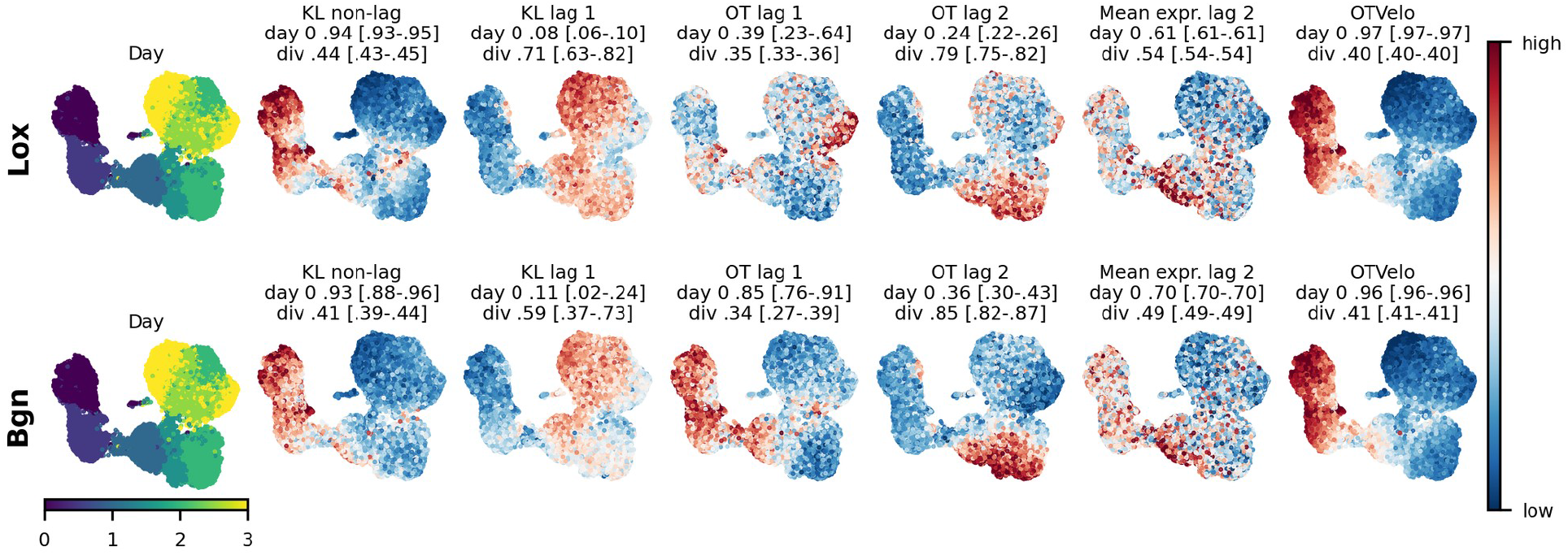
KL, OT, the mean-expression baseline and OTVelo [43] from the same query genes, Lox (top) and Bgn (bottom), on the reprogramming time course (UMAP of the cells of days 0–3), mean over the three training seeds. Columns: day, KL without lag and at lag 1, OT at lags 1 and 2, the mean-expression baseline at lag 2, and OTVelo, read forward from the query gene. Colour: Scanpy score genes of the top 30 retrieved genes, scaled for each seed and averaged over the seeds. Above each panel: AUROC of the score for the cells of day 0 and for the cells with a high cell-cycle signature (div: top quarter of the cell-cycle signature of Schiebinger et al. [31]), mean over the seeds with the range in brackets. The baseline and OTVelo do not depend on the seed. OT at lag 2 reaches the cells with a high cell-cycle signature from both query genes in all seeds, KL at lag 1 less consistently, and neither the baseline nor OTVelo reaches them. The values of each seed are in Table 19.

**Figure 3:**
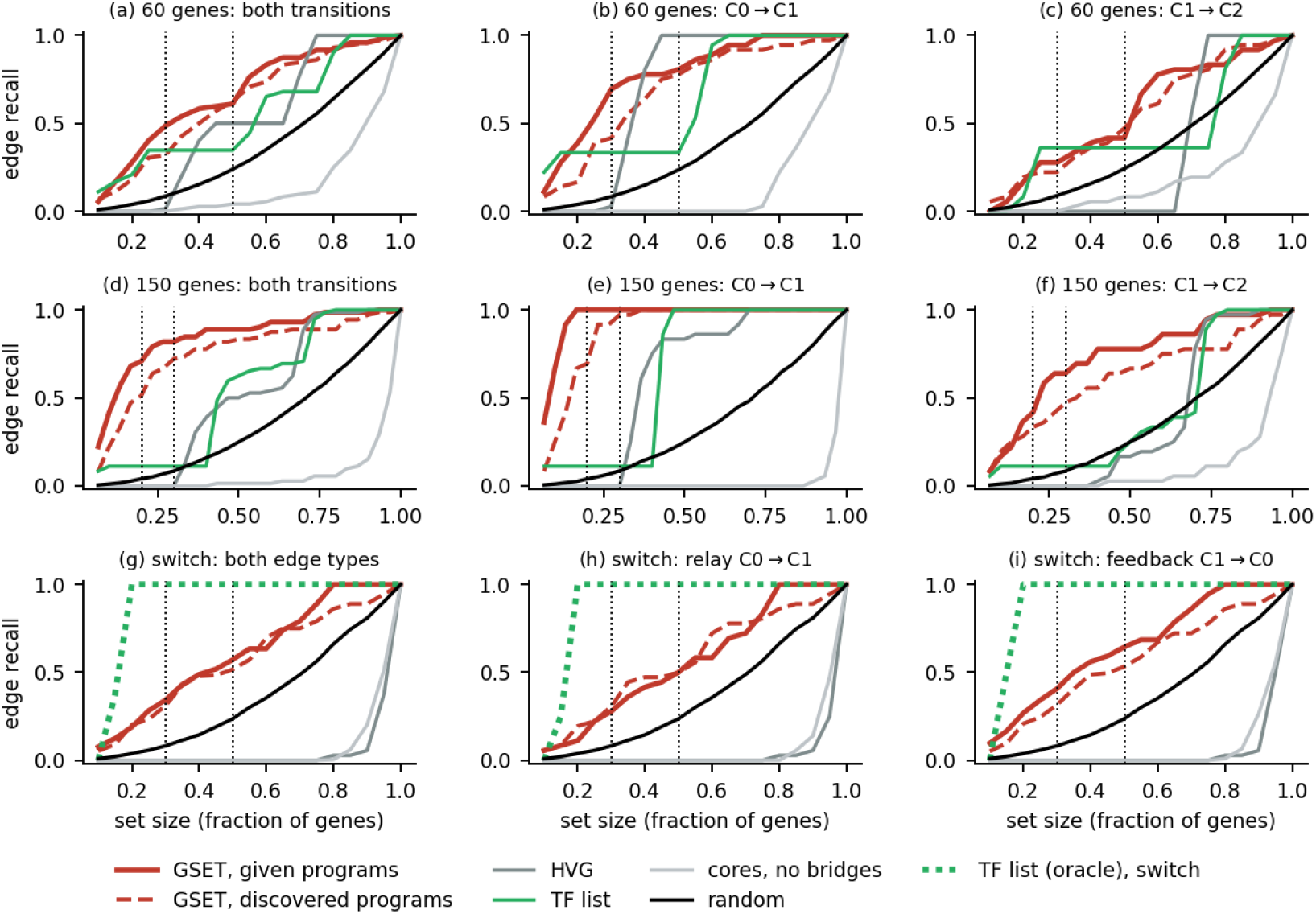
Retrieval of bridges: edge recall as a function of the size of the selected set. **a–c**: 60-gene cascades; **d–f**: 150-gene cascades (six networks each), for both transitions together (a, d), C0→C1 (b, e) and C1→C2 (c, f). **g–i**: switches (six networks), for both edge types together (g), the activating relay edges C0→C1 (h) and the feedback (inhibitory) edges C1→C0 (i); there all edges join TFs, so the TF list is an oracle (dotted green). GSET bridges with the programs given (solid) and discovered (dashed) are compared with HVG, the TF list, program cores and random subsets of the same size. Dotted vertical lines: budgets used in Table 2.

### 2.2 Gene embedding trajectories

#### Co-expression embedding

At each timepoint, GSET builds a WGCNA-style graph 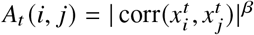 with a fixed soft-thresholding exponent *β* and encodes it with a two-layer graph convolutional network [15]. The node features are the genes’ correlation profiles, and the network is trained as a graph autoencoder with an inner-product decoder [14]. Its weights are shared across all timepoints, following weight tying in Siamese GNNs [17] and joint training in dynamic graph autoencoders [22]. This yields 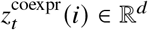 in which genes with similar co-expression context occupy compatible positions. Message passing averages a gene’s signal with its neighbours’, and correlation is scale-invariant, so this embedding does not directly encode each gene’s own expression level.

#### Marginal-expression embedding

To retain that information, a permutation-invariant DeepSets encoder [42], shared across genes and timepoints, maps gene *i*’s library-normalised expression values across the cells of timepoint *t* to 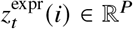. The encoder uses a per-gene standardisation that is fixed across timepoints, no message passing and no information from other genes. It is trained to reconstruct the empirical quantile function of the gene’s expression distribution, evaluated on a fixed grid of quantile levels, with an additional subsample-consistency term that penalises different embeddings for two random halves of the cells. Any representation of the expression distribution must be invariant to the order of the cells, since cells are unmatched across timepoints and vary in number. The consistency term encourages robustness to finite-cell sampling. Full construction is given in Appendix A. A learned embedding of the quantile function is used rather than the raw quantile vector, which also lets the encoder be changed without changing the downstream algorithm.

#### Signal-specific embedding spaces

The OT residual measures a gene’s deviation from a population-level shift, which is meaningful only if the space has a stable coordinate frame across timepoints and does not smooth over individual genes. The marginal-expression space provides both properties by construction. The co-expression space is defined only up to isometry across timepoints and is smoothed by message passing. Because the KL signal depends only on within-timepoint distances, it is well posed there. *δ*^OT^ is therefore computed in *z*^expr^ and *δ*^KL^ in *z*^coexpr^. Without the learned encoders, with the signals computed directly on the correlation profiles and the quantile vectors, the retrieval works as well or better on the 60-gene cascades and similar at a budget of 30 genes and somewhat worse at 45 on the 150-gene cascades, and its coverage drops sharply when 1000 unrelated genes are added (Appendices F and H), the regime of large gene panels that GSET targets.

### 2.3 OT residual: gene-specific deviation from the global shift

For a pair of timepoints (*s, t*), let 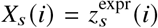 and define the empirical distribution 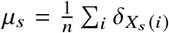, assigning equal mass to each gene. The global shift is estimated with entropy-regularised OT, solved by Sinkhorn iterations:

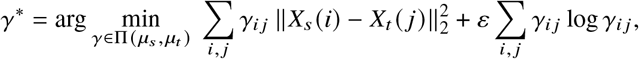

with a fixed regularisation strength *ε* = 0.05. The cost matrix is divided by its maximum, so *ε* is relative to the largest squared distance. The coupling is computed blind to gene identities. The OT-transported position of gene *i* is the barycentric projection 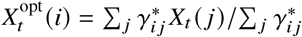, that is, the position predicted by the redistribution of the whole population. Only then is the gene correspondence reintroduced, and the residual of gene *i* is

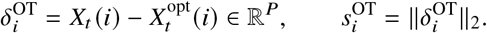

Unlike the raw displacement *X*_*t*_ (*i*) − *X*_*s*_ (*i*), the residual separates a gene’s own deviation from the collective movement. The OT plan is soft and is fitted to the population, so it can partly absorb a change that is aligned with the dominant shift, and very large coordinated programs may be partly attributed to the global trend. *δ*^OT^ should therefore be read as deviation from the population trend, not as total change.

### 2.4 KL signal: co-expression neighbourhood reorganisation

For gene *i* at timepoint *t*, a softmax distribution over the other genes is defined from squared distances in 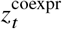:

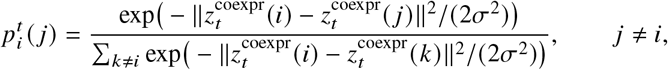

with *σ*^2^ set independently for each transition to the median of the non-zero squared distances between genes in 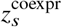. The scalar signal is the symmetrised divergence 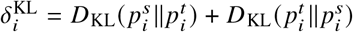. GSET also retains its directed per-partner contributions,

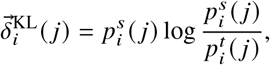

which record which relationships changed and are the input of the bridge retrieval of §2.6. Because 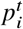 depends only on within-timepoint distances, *δ*^KL^ is invariant to translations and orthogonal transformations of either embedding.

### 2.5 Per-transition signals and program retrieval

Applying §2.3–2.4 to each consecutive pair (*t*_*τ*_, *t*_*τ*+1_), for *τ* = 0, …, *M* − 1, gives per-transition vectors 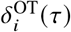 and 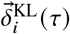. Initial timepoints at which the mean count over all genes is at most one, before the programs are expressed, are left out. The concatenations of these vectors over *τ* are *δ*_OT,concat_ and *δ*_KL,concat_. The reference representations without explicit differencing are *z*_global_, the co-expression embedding pooled over all timepoints, and *z*_concat_, the concatenation of the row-normalised 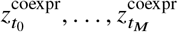.

Static program structure is recovered from cosine similarity in *z*_concat_, without supervision on program identity. For a query gene *q*, genes with high cosine similarity to *q* in *z*_concat_ define its static program context. Programs defined through the contexts of query genes can share genes, whereas programs found by clustering *z*_concat_ (§2.7) cannot, since each gene then belongs to exactly one program. In parallel, the concatenated KL and OT signals represent each gene by its trajectory of temporal change across transitions. Cosine similarity between these vectorial signals identifies genes undergoing coordinated temporal changes. These static and temporal representations are served interactively in Gene Program Explorer [40].

### 2.6 Lagged similarity and bridge retrieval

For a signal *r* ∈ {OT, KL} and a lag ℓ≥ 0, let

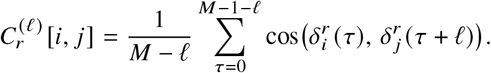

At ℓ= 0 this measures same-interval *co-change* and is symmetric. For ℓ≥ 1 it measures *statistical precedence*: the forward reading 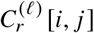 asks whether a change in *i* is followed ℓtransitions later by a similar change in *j*, whereas the reversed reading is 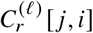. Which lag carries a temporal relationship depends on the sampling interval relative to the underlying dynamics (§3.4). GSET therefore treats combinations of signal, lag and direction as complementary *channels*, rather than assigning a fixed regulatory role to any single lag.

Bridge genes are retrieved between consecutive programs *C*_*a*_ → *C*_*b*_, in the order in which the programs become active (§2.7). For each channel *c*, a gene *i* ∈ *C*_*a*_ is scored by its best partner in the next program, 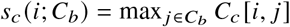, and a gene *j* ∈ *C*_*b*_ by its best partner in the previous one, 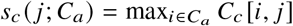, so that lagged channels are read forward. The co-change channel at lag 0 is symmetric and instead scores a gene by its best partner in any other program, *s*_*c*_ (*i*) = max_*j*∉*C* (*i*)_ *C*_*c*_ [*i, j*], where *C*(*i*) is the program of gene *i*. Each channel contributes a fixed number of genes to the budget. For a lagged channel, such as OT at lag 1, sources and targets are scored on different scales, by the best entry of a row and of a column of *C*_*c*_, and a relay edge is covered only when both are retrieved. Its share is filled alternately with the best remaining source and the best remaining target of each transition, each ranked separately (alternating allocation). The union across channels forms the bridge context *S*_*ab*_, and the union over consecutive program pairs forms *S*_bridge_. Taking the union rather than the intersection implements the preference for recall over precision stated in §1. Scoring genes rather than gene pairs makes the retrieval budget explicit in terms of the number of retained genes. The channel set is fixed globally and is never tuned per program pair. In the experiments of §3.6, it is {KL^(0)^, OT^(1)^ forward}, with the larger share of the gene budget given to KL^(0)^. This choice is motivated in §3.4.

Ground-truth edges are never used for ranking or selection.

### 2.7 Integration with mechanistic GRN inference

When program identities are unknown, they are discovered by agglomerative clustering of *z*_concat_ on the cosine distance, with the number of programs treated as known, and ordered by the half-rise time of the mean expression of each cluster. When they are given, their true activation order is used. Within each resulting cluster, program cores *S*_core_ are the top genes by within-cluster cosine centrality, combined with retrieved bridges into a fixed budget *S*_final_ = *S*_core_ ∪ *S*_bridge_. Cores give the downstream model the context of each program, that is, the genes that define it, so that a bridge gene is read relative to the programs it connects. They are a fixed share of the budget and are not selected to recover edges between programs. A dedicated GRN method (here CardamomOT [23]) is then fitted on *S*_final_. Lagged similarity can arise from shared upstream drivers as well as from direct regulation, so GSET only selects the context. Existence, direction and sign of edges are left to the mechanistic model.

## 3 Results

### 3.1 Datasets and baselines

#### Data

i. A Negative Binomial (NB) mixture with 40 genes and 5,000 cells per timepoint, in which co-expression rewiring and positional (marginal-expression) deviation are induced separately (Appendix B).
ii. HARISSA [8] mechanistic simulations with known ground-truth networks (design, sampling and number of runs in Appendix I). In a *cascade*, three gene programs C0→C1→C2 are activated in sequence, and consecutive programs are linked by *relay edges* from the TFs of one program to any gene of the next (TF-to-any relays), with 60 or 150 genes. In a *switch*, two mutually exclusive programs of 60 genes in total are activated one after the other. Two further families, in which every relay edge joins two TFs (150-gene cascades and a second kind of switch with edges between the programs), are used for retrieval and for CardamomOT (Appendix G). Unless stated otherwise, results are averaged over six networks per scenario and three training seeds of the encoders. Gene retrieval, the channel ablation and the CardamomOT experiments (§3.6) use a single seed.
iii. Mouse embryonic fibroblasts reprogrammed into induced pluripotent stem cells [31], serum condition, days 0–3: seven timepoints 12 h apart, 34,101 cells and 4,007 genes (Appendix K).
iv. A rhabdomyosarcoma (RMS) scRNA-seq dataset [30] (GEO: GSE248183) ordered by pseudotime into six bins (*t* = 0, 16, …, 80), 13,968 cells and 8,442 genes, with cells annotated as quiescent, intermediate or proliferative (Appendix L).

#### Baselines

*z*_global_ (static) and *z*_concat_ (trajectory of states), the raw displacement between two snapshots in the embedding space of each signal (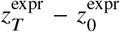 and 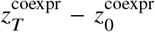), a lagged comparison of raw 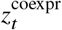 without differencing, OTVelo [43], and the OT gene–gene distance of Huizing et al. [11] that underlies GeneTrajectory [27] (“OT similarity”). The comparison uses this representation rather than the full pseudo-trajectory pipeline, which orders cells rather than selecting genes. For gene retrieval, the baselines also include the GSET selection procedure applied to lagged cosine similarities (lag 0 and lag 1 forward) of the centred mean-expression profiles (“lagged correlation”), DELVE [28], which uses neither time labels nor programs, the same procedure applied to the OTVelo and OT-similarity scores, and gene-level and network-based selections (temporal differential expression, co-expression hubs, Hotspot, GRNBoost2, GENIE3), all described in Appendix E. Not all baselines apply to every experiment below, and each subsection specifies which are compared.

### 3.2 OT and KL capture complementary forms of gene-level temporal change

On the NB-mixture dataset of §3.1, ranking genes by *δ*^KL^ and by *s*^OT^ reproduces the intended specialisation across two expression-level resolutions at low and moderate dropout. KL preferentially detects co-expression rewiring and the OT residual preferentially detects positional deviation (Appendix B, Figs. 4–5). The specialisation is relative rather than absolute, and the sensitivity of KL to rewiring decreases under strong dropout.

**Figure 4:**
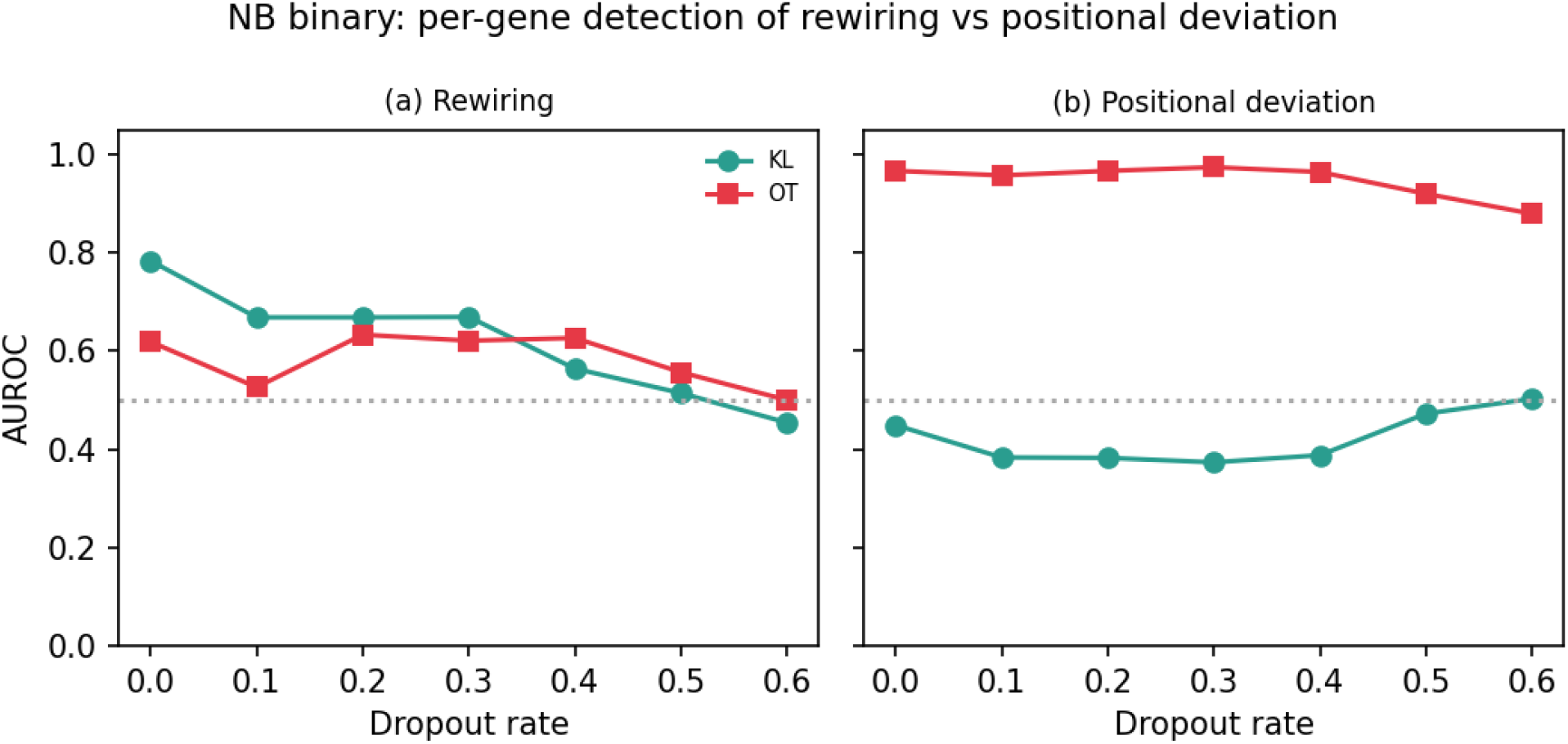
AUROC for detecting signal genes versus dropout in the binary NB benchmark. **(a)** Co-expression rewiring, with marginal expression held stable. **(b)** Positional deviation, with co-expression partners unchanged. Without dropout KL is more sensitive than OT to rewiring, and the OT residual is more sensitive than KL to positional deviation at all dropout rates. The sensitivity of KL to rewiring decreases as dropout increases. Curves show seed means.

**Figure 5:**
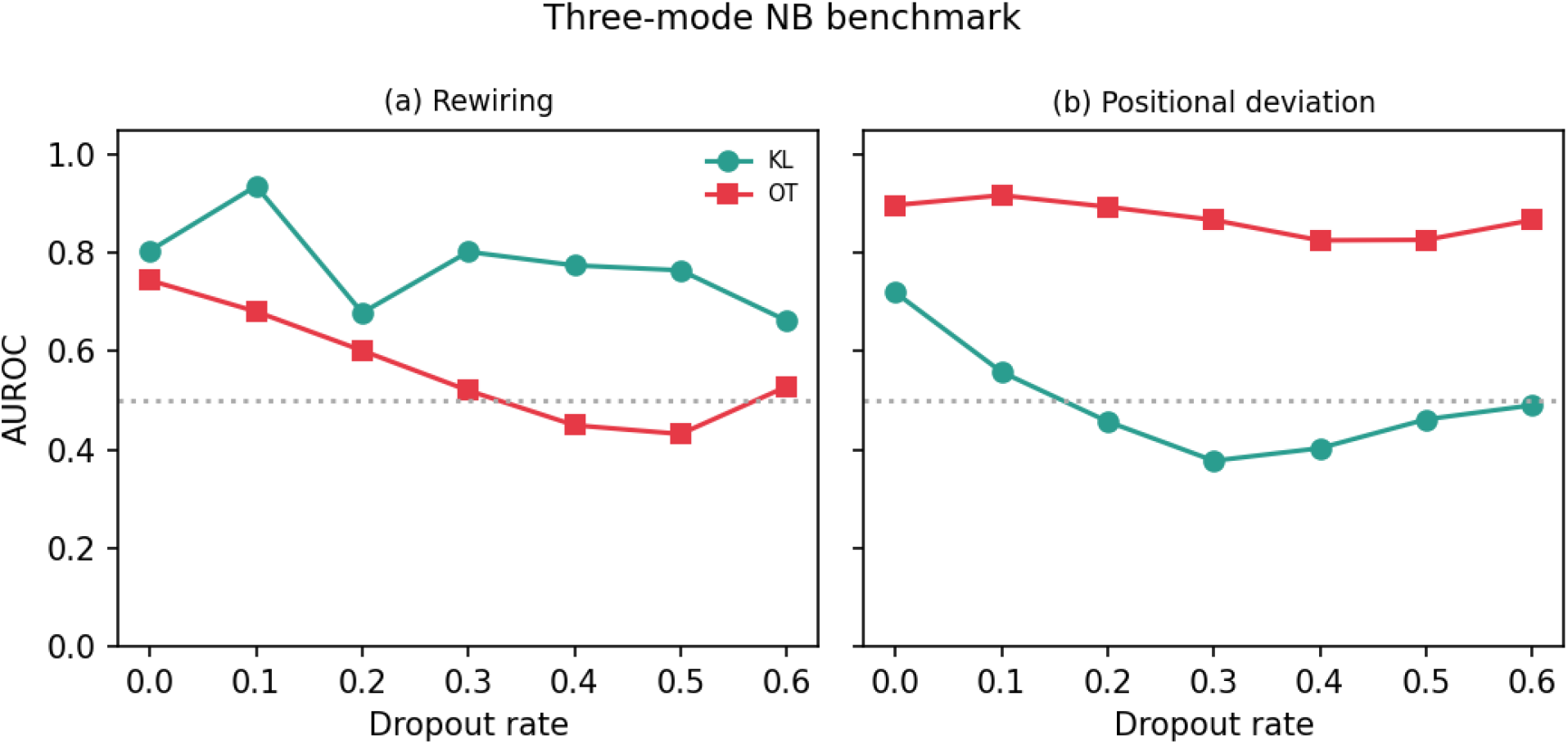
Extended three-level NB benchmark. **(a)** Rewiring: *δ*^KL^ detects the rewired anchors across dropout rates, and *s*^OT^ only at low dropout. **(b)** Positional deviation: *s*^OT^ gives the stronger signal and *δ*^KL^ is only weakly sensitive. Curves show seed means.

### 3.3 Temporal signals preserve program structure

The lagged similarity of §2.6 only makes sense if the per-transition signals carry program structure rather than noise. Each signal is therefore compared with naive differencing of the same embedding, on the first and last snapshots of a single run (Table 1, and Figs. 6–7 in Appendix C). Raw displacement loses much of this structure in the scenario that suits each signal, and the signals recover it. In the cascade, the within-minus between-program gap is 0.76 for the OT residual against 0.04 for 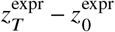. In the switch, it is 1.01 for the KL signal against 0.25 for 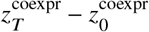. Program membership itself is best read from state representations, and *z*_concat_ and OT similarity are among the strongest in both scenarios (Table 3, Appendix C). This is why GSET defines programs from *z*_concat_ (§2.5). The signals are kept for a different purpose: they give one vector per transition, which says how each gene changed and when (for KL, also which of its relationships changed), and this is the input that the lagged similarity of §2.6 requires.

**Table 1.** Within-minus between-program cosine gap at the endpoints (single run, mean ± SD over six networks per scenario). Each signal is paired with its displacement baseline.

|  | Cascade | Switch |
|---|---|---|
| *z*<sub>*T*</sub><sup>coexpr</sup> − *z*<sub>0</sub><sup>coexpr</sup> (baseline) | 0.42 ± 0.14 | 0.25 ± 0.20 |
| KL vector | 0.36 ± 0.31 | 1.01 ± 0.78 |
| *z*<sub>*T*</sub><sup>expr</sup> − *z*<sub>0</sub><sup>expr</sup> (baseline) | 0.04 ± 0.05 | 1.25 ± 0.53 |
| OT residual | 0.76 ± 0.38 | 1.55 ± 0.46 |

**Table 2.** CardamomOT on retrieved contexts at a budget of 30 genes: relay-edge coverage per transition (of six; an uncovered edge counts as missed) and relay AUROC, which at budgets small relative to the gene panel is dominated by coverage, since uncovered pairs score 0, and therefore measures selection and inference together; all edges: AUROC over all true edges within the context for the model fitted on the context (ctx.) or on all genes restricted to the same pairs (full); chance 0.5. True/disc.: programs given/discovered. Program cores: cores without bridges. Lag-corr.: the same selection from lagged correlations of mean expression. DELVE: default parameters, without cores. Random: mean of three sets. Full: all genes. Further budgets and AUPRC: Table 7.

| Context | C0→C1 |  | C1→C2 |  | All edges |  |
|---|---|---|---|---|---|---|
|  | cov. | AUROC | cov. | AUROC | ctx. | full |
| *60-gene cascades (six networks)* |  |  |  |  |  |  |
| GSET (true) | 4.5 | 0.85 | 3.5 | 0.73 | 0.68 | 0.71 |
| GSET (disc.) | 3.8 | 0.78 | 3.0 | 0.70 | 0.67 | 0.70 |
| HVG | 6.0 | 0.95 | 0.0 | 0.50 | 0.66 | 0.66 |
| TF list | 2.0 | 0.61 | 2.2 | 0.65 | 0.65 | 0.65 |
| Program cores | 0.0 | 0.42 | 0.5 | 0.45 | 0.68 | 0.65 |
| Random | 1.5 | 0.52 | 1.5 | 0.52 | 0.69 | 0.68 |
| DELVE | 1.0 | 0.45 | 0.0 | 0.50 | 0.67 | 0.65 |
| Full | 6.0 | 0.88 | 6.0 | 0.86 | 0.68 | – |
| *150-gene cascades (six networks)* |  |  |  |  |  |  |
| GSET (true) | 5.5 | 0.93 | 1.3 | 0.59 | 0.70 | 0.65 |
| GSET (disc.) | 2.5 | 0.68 | 1.5 | 0.60 | 0.63 | 0.64 |
| HVG | 0.0 | 0.50 | 0.0 | 0.50 | 0.50 | 0.50 |
| TF list | 0.7 | 0.55 | 0.7 | 0.55 | 0.65 | 0.58 |
| Program cores | 0.0 | 0.49 | 0.0 | 0.49 | 0.65 | 0.64 |
| Random | 0.1 | 0.49 | 0.0 | 0.49 | 0.65 | 0.61 |
| DELVE | 0.0 | 0.50 | 0.0 | 0.50 | 0.67 | 0.65 |
| Lag-corr. (true) | 0.0 | 0.49 | 0.7 | 0.54 | 0.64 | 0.66 |
| Lag-corr. (disc.) | 1.7 | 0.63 | 0.8 | 0.56 | 0.66 | 0.61 |
| Full | 6.0 | 0.76 | 6.0 | 0.79 | 0.63 | – |

**Table 3.** Program recovery by Ward clustering of cosine-distance matrices, for all representations. Mean over six networks per scenario and three seeds. The last block contains ablations of the concatenations, not baselines.

|  | Switch |  |  | Cascade |  |  |
|---|---|---|---|---|---|---|
| Representation | ARI | NMI | AUROC | ARI | NMI | AUROC |
| *z*<sub>global</sub> | 0.97 | 0.95 | 0.99 | 0.06 | 0.16 | 0.87 |
| *z*<sub>concat</sub> | 1.00 | 1.00 | 1.00 | 0.74 | 0.77 | 0.89 |
| OT similarity | 1.00 | 1.00 | 1.00 | 0.76 | 0.80 | 0.96 |
| *δ*<sup>KL</sup> concat | 1.00 | 1.00 | 1.00 | 0.56 | 0.64 | 0.90 |
| *δ*<sup>OT</sup> concat | 0.57 | 0.60 | 0.79 | 0.73 | 0.76 | 0.94 |
| *δ*<sup>OT</sup> endpoint | 0.99 | 0.98 | 1.00 | 0.53 | 0.61 | 0.91 |
| *Ablations* |  |  |  |  |  |  |
| *z*<sub>concat</sub> + *δ*<sup>OT</sup> concat | 1.00 | 1.00 | 0.98 | 0.74 | 0.76 | 0.93 |
| *z*<sub>concat</sub> + *δ*<sup>KL</sup> concat | 1.00 | 1.00 | 1.00 | 0.68 | 0.72 | 0.90 |
| MaxRank(*δ*<sup>OT</sup> endpoint, *δ*<sup>OT</sup> concat) | 0.83 | 0.83 | 0.86 | 0.73 | 0.76 | 0.94 |
| MaxRank(*δ*<sup>OT</sup> concat, *δ*<sup>KL</sup> concat) | 0.74 | 0.76 | 0.86 | 0.76 | 0.79 | 0.93 |
| MaxRank(*δ*<sup>OT</sup> endpoint, *δ*<sup>KL</sup> concat) | 1.00 | 0.99 | 1.00 | 0.73 | 0.78 | 0.93 |

**Figure 6:**
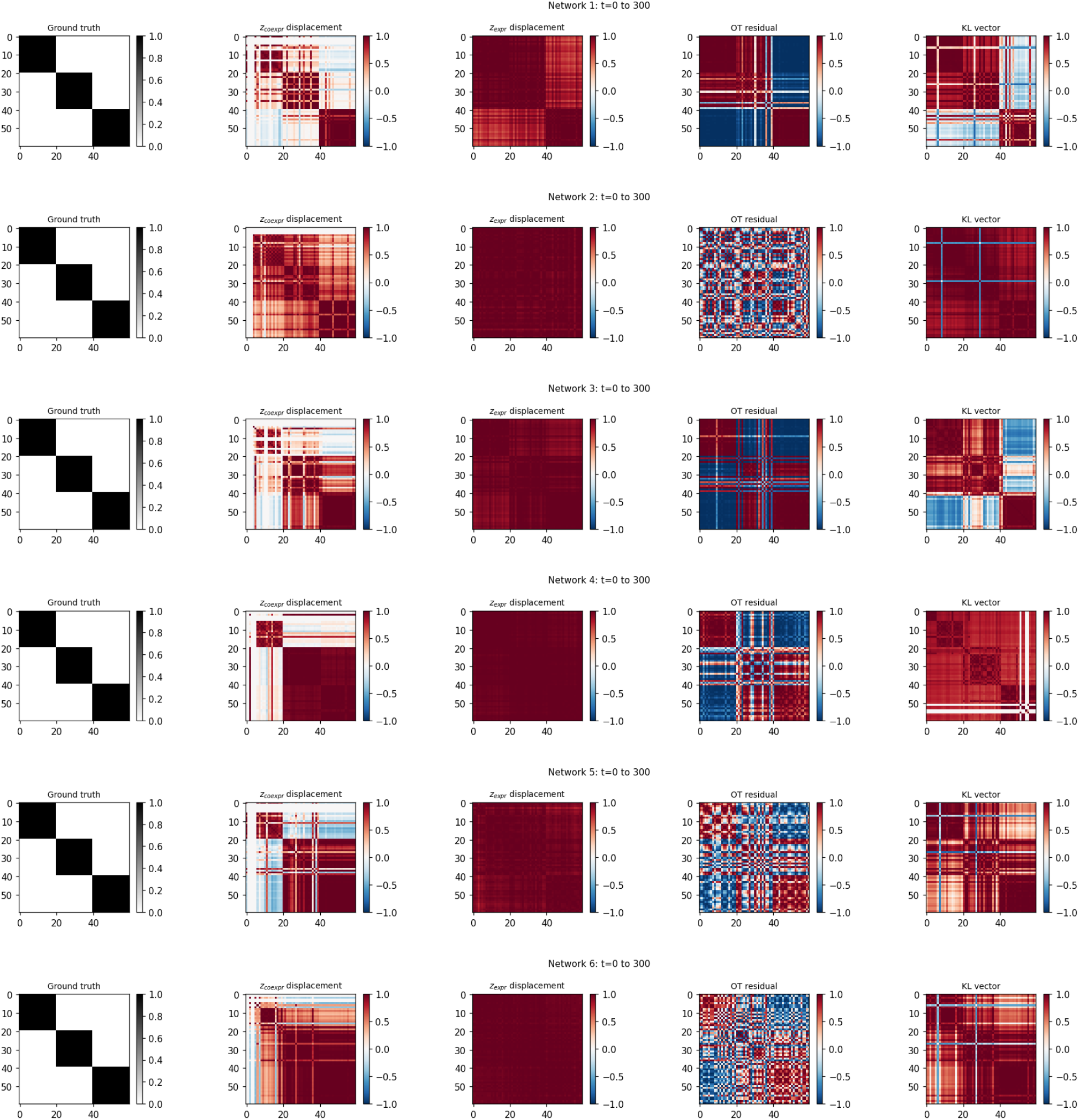
Cosine similarity of per-gene signal vectors at the endpoints in the six cascades (three programs of 20 genes), sorted by ground-truth program. Columns: ground truth (block-diagonal matrix of the true program labels); displacement 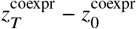; displacement 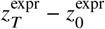; OT residual; KL vector. Rows: networks 1–6.

**Figure 7:**
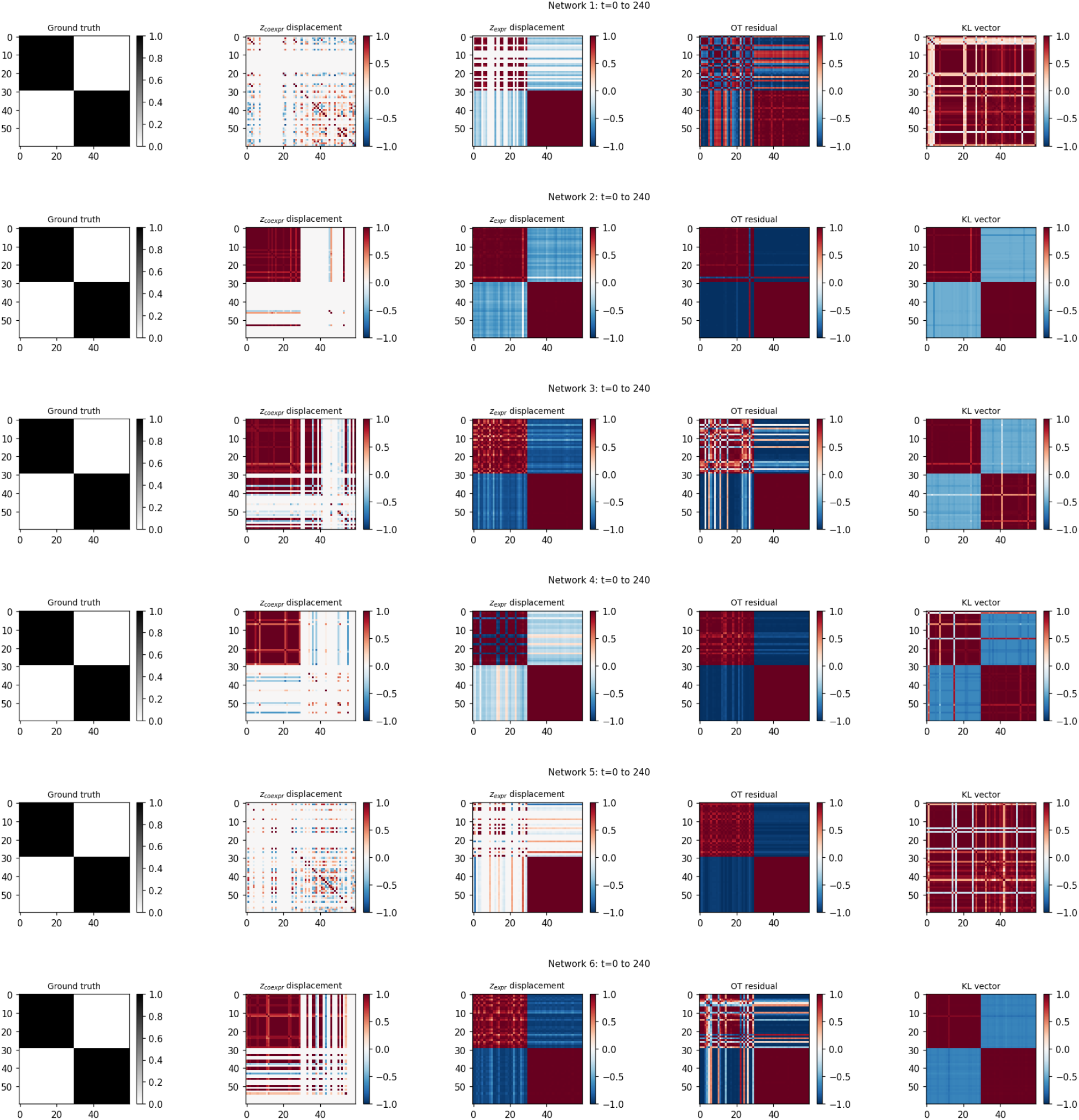
Cosine similarity of per-gene signal vectors at the endpoints in the six switches (two programs of 30 genes), sorted by ground-truth program. Columns and rows as in Fig. 6.

### 3.4 Lagged association between programs and its dependence on sampling resolution

Association between programs – whether the genes of one program change in a way that is related to the genes of the next – is distinct from the partition recovery of §3.3, since a representation can recover which genes belong together without identifying which relationships link programs. It is tested on both transitions (C0→C1, C1→C2) by the discrimination of true relay edges from non-edges matched on source/target program role and on the source being a TF (TF-to-gene pairs of the same two programs), measured by AUROC. Without this matching, a channel could score well merely by knowing program order or TF identity. Direction is reported as a secondary quantity, as *P*(*C*_*i j*_ > *C*_*ji*_) on true edges and as the separation of edges from non-edges by the asymmetry *C*_*i j*_ − *C*_*ji*_, since program order alone already gives a high value of the former. A channel can be informative for one transition without being informative for the other.

#### Edge discrimination

On the TF-to-any cascades (Table 4, Appendix D), both KL at lag 0 and OT at lag 1 discriminate relay edges on both transitions. KL at lag 0 is the strongest channel on C0→C1 and OT at lag 1 on C1→C2. Lagged KL channels beyond lag 0 are weaker.

**Table 4.**
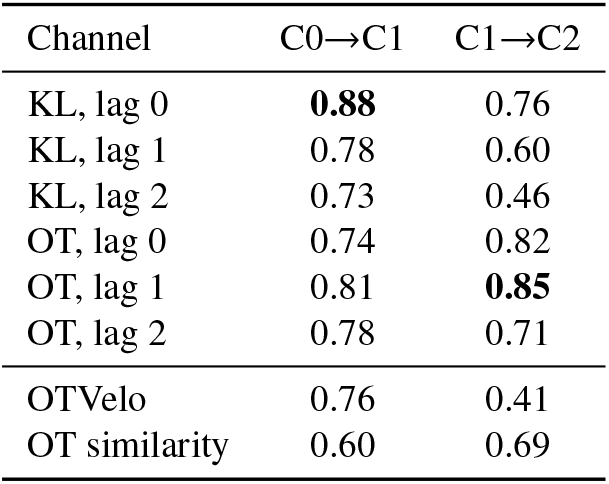
Discrimination of relay edges on the TF-to-any cascades (AUROC of true edges against all TF-to-gene pairs without an edge between the same two programs, forward reading; mean over six networks and three seeds). Chance is 0.5.

#### Direction

Program order alone already orients most pairs forward, so direction is informative only if it separates true relay edges from non-edges between the same programs. OT at lag 1 does so on both transitions: its asymmetry *C*_*i j*_ − *C*_*ji*_ is larger for relay edges than for non-edges (Table 5, Appendix D). OT at lag 2 does so more weakly on the second transition, and the KL channels hardly at all. The baselines, including OTVelo, separate them on at most one transition and fall to chance or below on the other, so their direction is not consistent across transitions.

**Table 5.** Direction on the TF-to-any cascades (mean over six networks and three seeds). *Edges*: fraction of true relay edges with *C*_*i j*_ > *C*_*ji*_. *Non-edges*: the same fraction among TF-to-gene pairs of the two programs without an edge. *AUROC*: separation of edges from these non-edges by the asymmetry *C*_*i j*_ − *C*_*ji*_ (0.5: no information beyond program order). Channels with symmetric similarity (lag 0, *z*_global_) have no direction.

|  | C0→C1 |  |  | C1→C2 |  |  |
|---|---|---|---|---|---|---|
| Channel | edges | non-edges | AUROC | edges | non-edges | AUROC |
| OT, lag 1 | 0.90 | 0.53 | **0.80** | 0.97 | 0.91 | **0.79** |
| OT, lag 2 | 0.92 | 0.65 | 0.79 | 0.97 | 0.90 | 0.65 |
| KL, lag 1 | 0.44 | 0.32 | 0.60 | 0.57 | 0.62 | 0.51 |
| KL, lag 2 | 0.31 | 0.23 | 0.59 | 0.61 | 0.68 | 0.50 |
| OTVelo | 1.00 | 0.94 | 0.70 | 1.00 | 1.00 | 0.18 |
| OT similarity | 0.86 | 0.84 | 0.48 | 0.19 | 0.06 | 0.77 |
| Lagged *z*<sub>*t*</sub><sup>coexpr</sup> | 0.12 | 0.08 | 0.77 | 0.88 | 0.87 | 0.55 |

#### The optimal lag follows sampling resolution

If co-change and precedence are the same relationship seen at different resolutions, the lag at which relay edges are best discriminated should shift as the sampling grid is refined (coarse, baseline and fine grids; Appendix I). This prediction holds for OT: its best lag grows as the grid is refined, from mostly lag 0 on the coarse grid to lag 1 on the baseline grid and lag 2 on the fine grid. For KL, lag 0 is the best on every grid, so its best lag does not shift (Table 6, Fig. 9, Appendix D). Because the optimal lag is not known in advance on experimental data, GSET uses within each signal the channel that stays informative on every grid: lag 0 for KL, which does not depend on the grid, and lag 1 for OT. These channels were fixed from this comparison, which uses only the 60-gene cascades, and not from any retrieval results. The 150-gene cascades of §3.6 were not used to choose them.

**Table 6.** Edge-discrimination AUROC by sampling resolution and lag (forward reading), averaged over the two transitions and three seeds, on the six TF-to-any cascades. The grids sample the same time course at a constant step of 60 time units (coarse, 6 snapshots), at the variable steps of the baseline grid (11 snapshots) and at a constant step of 8 time units (fine, 38 snapshots). Bold: best lag for the network and grid. Chance is 0.5; values below 0.5 mean that non-edges score higher than edges.

|  | Coarse |  |  | Baseline |  |  | Fine |  |  |
|---|---|---|---|---|---|---|---|---|---|
| Net | ℓ=0 | ℓ=1 | ℓ=2 | ℓ=0 | ℓ=1 | ℓ=2 | ℓ=0 | ℓ=1 | ℓ=2 |
| *KL* |  |  |  |  |  |  |  |  |  |
| s1 | **0.86** | 0.74 | 0.31 | **0.78** | 0.67 | 0.68 | **0.85** | 0.21 | 0.51 |
| s2 | **0.83** | 0.77 | 0.46 | **0.80** | 0.75 | 0.51 | **0.89** | 0.15 | 0.71 |
| s3 | **0.90** | 0.48 | 0.58 | **0.79** | 0.65 | 0.71 | **0.83** | 0.25 | 0.57 |
| s4 | **0.82** | 0.77 | 0.22 | **0.82** | 0.76 | 0.57 | **0.92** | 0.07 | 0.81 |
| s5 | **0.83** | 0.66 | 0.24 | **0.87** | 0.61 | 0.51 | **0.86** | 0.14 | 0.77 |
| s6 | **0.88** | 0.34 | 0.22 | **0.82** | 0.69 | 0.60 | **0.86** | 0.36 | 0.49 |
| *OT* |  |  |  |  |  |  |  |  |  |
| s1 | **0.85** | 0.71 | 0.46 | 0.82 | **0.84** | 0.79 | 0.72 | 0.74 | **0.77** |
| s2 | **0.82** | 0.81 | 0.63 | 0.77 | **0.82** | 0.71 | 0.77 | 0.82 | **0.84** |
| s3 | **0.88** | **0.88** | 0.58 | 0.81 | **0.83** | 0.73 | 0.68 | 0.74 | **0.76** |
| s4 | **0.87** | 0.83 | 0.55 | 0.84 | **0.90** | 0.77 | 0.61 | 0.68 | **0.74** |
| s5 | 0.79 | **0.81** | 0.62 | 0.71 | **0.79** | 0.71 | 0.46 | 0.50 | **0.62** |
| s6 | 0.71 | **0.79** | 0.65 | 0.72 | **0.80** | 0.75 | 0.57 | 0.66 | **0.71** |

### 3.5 GSET traces a transition between programs on real data

GSET was applied to a reprogramming time course of mouse embryonic fibroblasts (Data (iii)). Reprogramming begins with an early, proliferative phase in which cells lose fibroblast identity and cell-cycle programs rise in all cells [31], so the expected transition runs from the fibroblast program to the cell-cycle program. Starting from fibroblast genes (Fig. 2), GSET recovers this transition: without lag, KL retrieves the fibroblast program of day 0, and with lag, KL and OT retrieve genes of the cell-cycle program, expressed in the dividing cells of the following days. Two controls on the same data do not: neither a baseline built on the mean expression of the genes nor OTVelo [43] reaches the cell-cycle program. The result depends on the query gene: of ten extracellular-matrix genes tested as queries, lagged KL separates the dividing cells from the others (AUROC above 0.5) for eight and lagged OT for three (Appendix K).

OT reaches the cell-cycle program at lag 2, not at lag 1, and this is in line with §3.4. In the simulations, the best OT lag increased as sampling became denser relative to the delay between programs, from lag 1 on the baseline grid to lag 2 on the fine grid. Here, the rise of the cell-cycle program follows the loss of fibroblast identity over several half-day steps. The lag was not chosen in advance, so the observation is consistent with the resolution effect of §3.4 but does not test it independently. It does show why GSET handles lags as separate channels (§2.6). On a new grid, adding OT at lag 2 to the union costs a few genes of budget, whereas leaving it out may miss the bridge genes of that transition.

Between the fibroblast and the cell-cycle programs, GSET retrieves bridge genes that are reproducible across training seeds, with about 50 of 200 shared between pairs of seeds against about 10 expected by chance (Appendix K). They include subunits of SAGA, a chromatin-modifying complex required at the onset of reprogramming [9], the early, proliferative phase in which GSET detects the transition. The data provide no ground truth, so these bridges remain candidates. GSET thus narrows thousands of genes to a few hundred candidates associated with the transition, which a mechanistic model and perturbation experiments can then test.

On rhabdomyosarcoma data ordered by pseudotime (Data (iv)), lagged KL similarly traces the progression from quiescent to intermediate and proliferative cells, starting from some genes of quiescent cells and in a query-dependent way (Appendix L).

### 3.6 GSET as a retrieval layer for mechanistic GRN inference

GSET acts as a retrieval layer upstream of CardamomOT (§2.7): it selects a compact gene context, so that the cost of the downstream model is set by the retrieval budget rather than by the number of genes (§1, Appendix J). It is compared at equal gene budget with selections of the same size: the most variable genes (HVG), a list of the TFs, the program cores without bridges, random subsets, DELVE and the lagged-correlation selection (Fig. 3, Table 2; switches and TF-to-TF relays in Appendix G). The number of program cores is fixed, and the budget is varied through the bridges.

The ground truth is the set of relay edges between consecutive programs, and a relay edge is *covered* if both its endpoints are retrieved. Programs are either *given*, with the true labels and activation order, or *discovered* without labels (§2.7). CardamomOT networks are scored by AUROC and AUPRC on the relay edges, an uncovered edge counting as missed, and on all edges within the retrieved set (Appendix E).

#### Retrieval

GSET bridges cover more relay edges than the program cores and random sets on every cascade, and more than HVG and the TF list on the 150-gene cascades (Fig. 3). Static selections miss these edges for structural reasons: HVG fills the set with the genes of the first program, the program cores are the most central genes of their programs (§2.7) while the endpoints of relay edges are less central, and the TF list contains the sources of relay edges but few of their targets. On the 150-gene cascades, where the targets are rarely TFs, GSET covers most relay edges of the first transition with 20% of the genes, whereas the static selections cover almost none. On the 60-gene cascades, HVG covers all relay edges of the first transition at a budget of 30 genes, but almost none at a budget of 18. The TF list does better only at the smallest budgets on the 60-gene cascades and when every relay edge joins two TFs, so that it acts as an oracle (Appendix G). Most of the coverage comes from KL at lag 0, and OT at lag 1 adds the edges of the second transition that KL covers least. With the programs given, the two channels together cover more relay edges of the second transition than either channel alone at the larger budget on the 150-gene cascades (Appendix F). The other dynamic and network-based selections, among them DELVE, OTVelo and GRNBoost2, also cover fewer relay edges than GSET on the 150-gene cascades (Appendix E). The advantage persists when 1000 real genes unrelated to the network are added to the 150-gene cascades. GSET then covers fewer relay edges than without them, but still more than the static selections at budgets of 30 and 45 genes (Appendix H).

#### CardamomOT on the retrieved contexts

Better coverage translates into better inference (Table 2). With the programs given and a budget of 30 genes, CardamomOT recovers the relay edges better on the GSET context than on the program cores, random sets and the TF list on every cascade. HVG is better only on the first transition of the 60-gene cascades and recovers no edge of the second. On the 150-gene cascades, where the static selections recover almost no relay edge, the GSET context recovers those of the first transition and some of the second. With discovered programs the advantage is smaller but remains, and at the smallest budget the TF list is comparable or better (Table 7). On the second transition the full-gene model remains clearly better than any retrieved context.

**Table 7.** CardamomOT on retrieved contexts, all budgets, with AUPRC/base. Columns as in Table 2, with AUPRC/base added for the relay edges of each transition and for all edges within the context (model fitted on the context). Chance: AUROC 0.5, AUPRC/base 1 (relay-edge base rate 0.015 for 60 genes, 0.0024 for 150). *60 genes*: six cascades with TF-to-any relays; six cores, the other genes are bridges (selected by KL at lag 0 and OT at lag 1 in a ratio of 2 to 1, with the allocation of §2.6 for OT). *150 genes*: six cascades with TF-to-any relays; budget 30: 10 cores and 20 bridges (14 selected by KL at lag 0 and 6 by OT at lag 1), budget 45: 15 cores and 30 bridges (20 by KL and 10 by OT). Rows as in Table 2. For 60 genes the coverage of Random is the expected value. TF list: annotated TFs filled up to the budget with HVG. GSET (KL only), DELVE, OTVelo, OT similarity and GRNBoost2 in GSET (the GSET procedure with GRNBoost2 edge weights, all genes as candidate regulators) are the further baselines described below, run at a budget of 30 genes only.

|  | C0→C1 |  |  | C1→C2 |  |  | All edges |  |  |
|---|---|---|---|---|---|---|---|---|---|
| Context | cov. | AUROC | AUPRC/b | cov. | AUROC | AUPRC/b | ctx. | full | AUPRC/b |
| *60 genes, budget 18* |  |  |  |  |  |  |  |  |  |
| GSET (true) | 2.3 | 0.67 | 20.3 | 1.5 | 0.61 | 8.0 | 0.72 | 0.72 | 2.3 |
| GSET (disc.) | 1.0 | 0.56 | 8.3 | 1.3 | 0.59 | 4.6 | 0.66 | 0.71 | 1.8 |
| HVG | 0.2 | 0.51 | 1.6 | 0.0 | 0.50 | 1.0 | 0.60 | 0.58 | 1.2 |
| TF list | 2.0 | 0.65 | 16.9 | 2.2 | 0.64 | 11.4 | 0.68 | 0.67 | 2.3 |
| Program cores | 0.0 | 0.47 | 1.0 | 0.0 | 0.46 | 1.0 | 0.75 | 0.63 | 2.6 |
| Random | 0.5 | 0.51 | 4.5 | 0.5 | 0.49 | 2.5 | 0.70 | 0.67 | 2.2 |
| *60 genes, budget 30* |  |  |  |  |  |  |  |  |  |
| GSET (true) | 4.5 | 0.85 | 30.9 | 3.5 | 0.73 | 14.6 | 0.68 | 0.71 | 1.9 |
| GSET (disc.) | 3.8 | 0.78 | 24.0 | 3.0 | 0.70 | 16.2 | 0.67 | 0.70 | 1.8 |
| HVG | 6.0 | 0.95 | 36.3 | 0.0 | 0.50 | 1.0 | 0.66 | 0.66 | 1.6 |
| TF list | 2.0 | 0.61 | 12.5 | 2.2 | 0.65 | 13.7 | 0.65 | 0.65 | 1.7 |
| Program cores | 0.0 | 0.42 | 1.0 | 0.5 | 0.45 | 2.9 | 0.68 | 0.65 | 2.1 |
| Random | 1.5 | 0.52 | 8.0 | 1.5 | 0.52 | 6.9 | 0.69 | 0.68 | 2.0 |
| GSET (KL only, true) | 4.8 | 0.88 | 38.7 | 2.5 | 0.66 | 11.8 | 0.67 | 0.71 | 1.8 |
| GSET (KL only, disc.) | 3.5 | 0.75 | 26.5 | 2.7 | 0.63 | 8.2 | 0.64 | 0.69 | 1.6 |
| DELVE | 1.0 | 0.45 | 7.1 | 0.0 | 0.50 | 1.0 | 0.67 | 0.65 | 1.7 |
| OTVelo (true) | 0.7 | 0.48 | 5.5 | 0.3 | 0.45 | 1.4 | 0.70 | 0.69 | 2.0 |
| OTVelo (disc.) | 1.2 | 0.53 | 7.9 | 2.2 | 0.62 | 7.2 | 0.66 | 0.68 | 1.8 |
| OT similarity (true) | 2.8 | 0.69 | 18.5 | 2.7 | 0.65 | 13.8 | 0.70 | 0.70 | 2.0 |
| OT similarity (disc.) | 1.2 | 0.52 | 12.1 | 1.2 | 0.52 | 3.5 | 0.67 | 0.67 | 1.9 |
| GRNBoost2 in GSET (true) | 0.0 | 0.42 | 1.0 | 4.2 | 0.77 | 16.6 | 0.69 | 0.68 | 1.8 |
| GRNBoost2 in GSET (disc.) | 1.0 | 0.50 | 3.7 | 2.3 | 0.60 | 12.8 | 0.67 | 0.69 | 1.8 |
| Full (60 genes) | 6.0 | 0.88 | 21.2 | 6.0 | 0.86 | 19.1 | 0.68 | – | 1.9 |
| *150 genes, budget 30* |  |  |  |  |  |  |  |  |  |
| GSET (true) | 5.5 | 0.93 | 217 | 1.3 | 0.59 | 66.8 | 0.70 | 0.65 | 3.0 |
| GSET (disc.) | 2.5 | 0.68 | 77.9 | 1.5 | 0.60 | 18.5 | 0.63 | 0.64 | 2.2 |
| HVG | 0.0 | 0.50 | 1.0 | 0.0 | 0.50 | 1.0 | 0.50 | 0.50 | 1.0 |
| TF list | 0.7 | 0.55 | 13.8 | 0.7 | 0.55 | 24.0 | 0.65 | 0.58 | 2.5 |
| Program cores | 0.0 | 0.49 | 1.0 | 0.0 | 0.49 | 1.0 | 0.65 | 0.64 | 2.9 |
| Random | 0.1 | 0.49 | 4.8 | 0.0 | 0.49 | 1.0 | 0.65 | 0.61 | 2.8 |
| Lag-corr. (true) | 0.0 | 0.49 | 1.0 | 0.7 | 0.54 | 32.1 | 0.64 | 0.66 | 2.2 |
| Lag-corr. (disc.) | 1.7 | 0.63 | 73.3 | 0.8 | 0.56 | 27.1 | 0.66 | 0.61 | 2.6 |
| GSET (KL only, true) | 5.3 | 0.94 | 294 | 1.0 | 0.56 | 33.4 | 0.66 | 0.63 | 2.5 |
| GSET (KL only, disc.) | 3.8 | 0.79 | 147 | 1.3 | 0.60 | 31.6 | 0.67 | 0.63 | 2.8 |
| DELVE | 0.0 | 0.50 | 1.0 | 0.0 | 0.50 | 1.0 | 0.67 | 0.65 | 2.7 |
| OTVelo (true) | 0.5 | 0.53 | 20.7 | 0.0 | 0.49 | 1.0 | 0.66 | 0.64 | 2.3 |
| OTVelo (disc.) | 1.2 | 0.59 | 40.7 | 0.5 | 0.53 | 18.6 | 0.64 | 0.64 | 2.5 |
| OT similarity (true) | 1.2 | 0.59 | 61.0 | 0.2 | 0.50 | 12.5 | 0.63 | 0.67 | 1.9 |
| OT similarity (disc.) | 1.0 | 0.57 | 53.0 | 0.2 | 0.51 | 12.5 | 0.60 | 0.68 | 2.0 |
| GRNBoost2 in GSET (true) | 0.0 | 0.49 | 1.0 | 1.0 | 0.56 | 42.0 | 0.65 | 0.63 | 2.5 |
| GRNBoost2 in GSET (disc.) | 1.5 | 0.60 | 35.6 | 0.8 | 0.56 | 29.1 | 0.67 | 0.70 | 2.5 |
| *150 genes, budget 45* |  |  |  |  |  |  |  |  |  |
| GSET (true) | 6.0 | 0.98 | 267 | 3.2 | 0.75 | 78.1 | 0.66 | 0.63 | 2.7 |
| GSET (disc.) | 4.2 | 0.82 | 124 | 1.8 | 0.62 | 52.2 | 0.66 | 0.63 | 2.7 |
| HVG | 0.0 | 0.50 | 1.0 | 0.0 | 0.50 | 1.0 | 0.51 | 0.49 | 1.0 |
| TF list | 0.7 | 0.54 | 7.3 | 0.7 | 0.55 | 21.5 | 0.58 | 0.58 | 2.1 |
| Program cores | 0.0 | 0.48 | 1.0 | 0.0 | 0.48 | 1.0 | 0.60 | 0.59 | 1.9 |
| Random | 0.4 | 0.52 | 16.8 | 0.9 | 0.55 | 15.3 | 0.64 | 0.62 | 2.4 |
| Lag-corr. (true) | 0.2 | 0.49 | 2.0 | 1.2 | 0.55 | 25.6 | 0.64 | 0.64 | 2.0 |
| Lag-corr. (disc.) | 1.8 | 0.57 | 41.2 | 0.8 | 0.55 | 22.1 | 0.63 | 0.59 | 2.4 |
| Full (150 genes) | 6.0 | 0.76 | 115 | 6.0 | 0.79 | 40.0 | 0.63 | – | 2.4 |

#### All edges

Scored on every true edge within the retrieved context, the model fitted on the GSET context recovers these edges about as well as the model fitted on all genes and restricted to the same gene pairs, as do most of the other selections (last two columns of Table 2). The contexts differ in their gene pairs, so the selections are not compared with each other on this measure, which is read as *no loss* from compression rather than as a gain.

## 4 Discussion

GSET separates program *identity* from program *dynamics*. Temporal embedding similarity (*z*_concat_) is among the strongest representations of which genes belong together. GSET’s vectorial signals add what that representation lacks: which relationships change, and in which transition. KL tracks co-expression rewiring and OT tracks deviations in marginal expression. Their specialisation is relative, and both detect the same program when a change has both components, as at the endpoints of the switch (Appendix C).

The association experiments support two conclusions. First, the order in which programs change is not by itself evidence of regulation: program order already orients most pairs forward, and most channels do not separate relay edges from non-edges by their *direction* consistently across transitions, and only the OT channels, most clearly at lag 1, do so on both (§3.4). Second, the lag that carries a relationship shifts with the sampling grid (§3.4), so neither co-change nor any single positive lag reliably indicates the regulatory role of a gene pair. Co-change channels are also needed in their own right, because a single regulator can drive simultaneous effects across several programs rather than acting through one program to reach the next [25]. The two signals therefore play complementary roles: KL at lag 0 carries most of the bridge retrieval, and OT at lag 1 adds relay edges of the second transition when the programs are given, as well as the most consistent direction information beyond program order (Appendix F).

Static selections work only when the relay edges happen to join the genes they favour. When the selected set is a small share of the genes and the targets of the relay edges are not TFs, they cover almost none of these edges, whereas GSET still covers a large part of them. The selection therefore matters most when the gene panel is small relative to the transcriptome, which is exactly when the mechanistic model is expensive. On real reprogramming data, lagged KL and OT trace the expected transition from the fibroblast to the cell-cycle program, which the mean-expression baseline and OTVelo do not (§3.5).

### Limitations and outlook

Three limitations bound these claims. First, lagged similarity is an association between programs and not causality. Bridge genes are candidates, which curated priors such as OmniPath [36] or DoRothEA [7] could filter further, and the existence, direction and sign of edges are left to the mechanistic model. Second, the validation has a limited scope. The ground truth comes from synthetic networks of at most 150 genes, because simulating larger networks with meaningful dynamics is itself difficult. What matters for the advantage of retrieval is likely not network size as such but the fraction of genes covered by the budget and whether the targets of relay edges are TFs, which decides whether a TF list can cover both endpoints, and the gap with static selections already widens from 60 to 150 genes. These networks have programs activated in sequence and linked by relay edges, the setting GSET is designed for. Networks with feedback between programs, overlapping programs or several parallel paths without a clear activation order are covered only partly, by the switches. Adding 1000 real genes unrelated to the network reduces the coverage of relay edges, although GSET still covers clearly more of them than the static selections (Appendix H). Replacing the cosine similarity of KL at lag 0 by its mutual rank, an exploratory variant, partly restores the coverage, so making the retrieval robust to many unrelated genes is a next step. On real data, the bridges retrieved from the reprogramming time course have no ground truth, and the RMS pseudotime partly encodes the progression it is used to read (Appendix L). Third, some components remain open. Programs are discovered with their number given, and the advantage is then smaller than with the programs given. On the reprogramming data they are defined around query genes, which needs no clustering and no number of clusters, only one query gene per program (§3.5). OT at lag 1 adds relay edges of the second transition, but not in every setting (Appendix F), and its best lag depends on the sampling grid (§3.4, §3.5). Both vectorial signals weaken under strong dropout, and GSET was evaluated upstream of a single mechanistic method, CardamomOT. These limitations define the next steps: estimating the number of programs and the OT lag from the data, and testing GSET upstream of other mechanistic methods. GSET narrows the transcriptome to a context in which candidate regulators of a transition can be sought. Turning these candidates into directed, signed edges is the role of a mechanistic model such as CardamomOT, and showing that they control cell-state transitions will require perturbation experiments, which this work prepares but does not establish.

## Code availability

Source code with documentation: https://anonymous.4open.science/r/gset-recomb/. The package is available on PyPI as gset (https://pypi.org/project/gset/). Gene Program Explorer: https://rmsexplorer.org, archived at Zenodo [40].

## A Gene embedding trajectories: full construction

Hyperparameters (*β* = 6, *τ* = 0.01, *L* = 2, network widths, *λ*_cons_ = 1) are given in the code repository.

### Co-expression embedding

Let *G* = {*g*_1_, …, *g*_*n*_} be genes observed at *t*_0_ < · · · < *t*_*M*_. At each timepoint, the Pearson correlation matrix 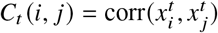 defines a weighted graph with soft-thresholded weights *A*_*t*_ ∈ [0, 1]^*n*×*n*^ [19]. A GCN [15] with weights shared across timepoints takes *H*^(0)^ = *C*_*t*_ and propagates

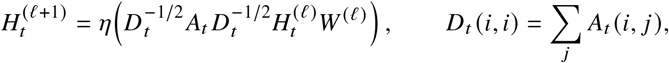

where *η* is the ELU nonlinearity for the first layer and the identity for the last, and no additional self-loops are added. Weights below *τ* are discarded. If no pair of genes passes the threshold at a timepoint, the graph is built from the strongest one percent of the gene pairs instead. The final layer gives 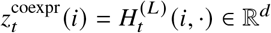. The encoder is trained as a graph autoencoder [14] with a raw inner-product decoder 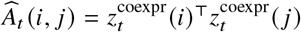 and loss

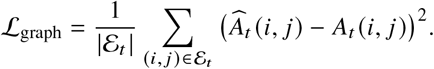

The experiments on real data (Appendices K and L) use a variant of this encoder for panels of thousands of genes, whose node features are the 50 leading singular vectors of the correlation matrix, with self-loops, a hidden width of 128 and an embedding of dimension 32 (code repository).

### Marginal-expression embedding

For gene *i* at timepoint *t*, let 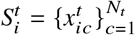 be its library-normalised log-expression values. Values are standardised with a fixed reference, 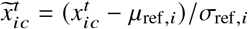, where *µ*_ref,*i*_, *σ*_ref,*i*_ are the mean and standard deviation of gene *i* over the training matrices and are the same for all timepoints. A DeepSets encoder [42] maps each value through 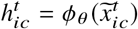 aggregates 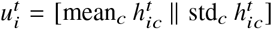, and outputs 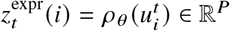. A decoder *D*_*θ*_ reconstructs the empirical quantile vector 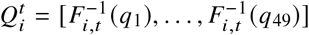, with *q*_*k*_ equally spaced in [0.01, 0.99], through

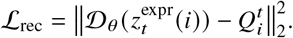

Each quantile level of the target is standardised across the genes of the same matrix before reconstruction. For two random halves of the cells of the same gene and timepoint, a consistency term penalises differing embeddings:

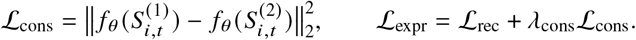

## B Negative Binomial mixture simulation: design and extended benchmark

This appendix details the synthetic benchmark of §3.2. AUROC compares signal genes with all remaining genes rather than with a matched null distribution. Under finite sampling, dropout, or coupling between expression distributions and measured correlations, the non-targeted signal therefore need not sit exactly at 0.5. Results are interpreted through the relative specialisation of the two signals.

Snapshots are simulated from a Negative Binomial mixture over *G* = 40 genes and *C* = 5,000 cells per timepoint, with *K* = 4 latent cell populations. Counts are drawn from NB(*r* = 2, *µ*_*k*_ (*g*)) with *µ*_high_ = 8.0 and *µ*_low_ = 0.5, using a split-cluster design:

| Population | Cells | High-expression genes |
|---|---|---|
| 0*a* | $\frac{1}{4}C$ | genes 0–6 |
| 0*b* | $\frac{1}{4}C$ | genes 7–13 |
| 1 | $\frac{1}{4}C$ | genes 14–26 |
| 2 | $\frac{1}{4}C$ | genes 27–39 |

Populations 0*a* and 0*b* play equivalent roles but contain disjoint high-expression genes. Genes can therefore exchange co-expression partners between the two populations while keeping their own marginal expression level, which gives a controlled rewiring task with no positional change.

Both embeddings are built as in Appendix A, with 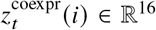. *δ*^KL^ and *s*^OT^ are computed as in §2.3–2.4. Dropout is applied independently to each count with probability *p*_*d*_ ∈ [0, 0.6].

### Rewiring task

Genes 0–3 are anchors and remain highly expressed in population 0*a* at both timepoints, so their marginal expression is unchanged. Genes 4–6 move from 0*a* to 0*b*, and genes 7–10 move from 0*b* to 0*a*. Each anchor therefore loses one set of partners and gains another. The four anchors are the signal genes.

### Positional-deviation task

The mean expression of genes 14–21 is multiplied by 3 in every population. Their on/off pattern is preserved while their marginal distributions shift, so the perturbation is intended to leave the co-expression partners unchanged. These eight genes are the signal set.

Performance is the AUROC for separating signal genes from all remaining genes, averaged over 10 simulation draws per dropout level and 5 training seeds. In each seed the GCN is trained on 30 matrices and the DeepSets encoder on 15 matrices, drawn from the unperturbed mixture, and the scores are evaluated on new draws. Without dropout the specialisation is clear: KL is higher than OT for rewiring, and OT is much higher than KL for positional deviation. As dropout increases, correlations between genes weaken, and at any nonzero dropout in the binary setting no gene pair passes the threshold, so the graph is built from the strongest pairs (Appendix A). In this regime the KL signal gradually loses its sensitivity to rewiring and reaches chance level at the highest dropout, whereas the OT score for positional deviation stays high. In the positional task the KL score is not at chance but mostly below it, which means that shifted genes receive lower KL scores than the other genes. A change of expression level also alters the estimated correlations, so the KL signal is not completely blind to it (Fig. 4).

### Three-level extension

The binary states are replaced by three on-levels, *µ* = 4, 10, 18, and an off-level *µ* = 0.3, keeping 0*a* and 0*b* role-equivalent. For rewiring, the anchors are the high-expression genes of the pair (genes 5, 6 in 0*a* and 12, 13 in 0*b*). Genes 0–2 move from 0*a* to 0*b* and genes 7–9 from 0*b* to 0*a*. For positional deviation, the mean expression of genes 14–21 is multiplied by 1.5 in every population. A factor of 3 at these levels changes cell composition enough to perturb co-expression, so the task would no longer be approximately positional. The same specialisation holds (Fig. 5). *δ*^KL^ detects the rewired anchors across dropout rates, with the highest values at low dropout and lower values under strong dropout, whereas *s*^OT^ is above chance at low dropout and falls to chance from moderate dropout. In this setting the graph is also built from the strongest pairs from a dropout rate of 0.3 onwards. In the positional task *s*^OT^ is stronger and stable across dropout, whereas *δ*^KL^ decreases from a moderate value without dropout to chance level.

## C Program recovery: endpoint maps, robustness to dropout and all representations

Figures 6 and 7 show, for each of the six networks of the two HARISSA [8] scenarios of §3.1, the cosine similarity between the signal vectors of all pairs of genes, with genes sorted by ground-truth program. Each signal is computed between the first and the last snapshot and is compared with its displacement baseline (§3.1). A displacement shows that a gene’s latent representation moved, but its coordinates are latent dimensions and do not decompose the change into contributions of specific relationships, unlike *δ*➔^KL^. The gaps between within-program and between-program similarity are given in Table 1.

In the cascade (Fig. 6), the OT residual shows block structure aligned with the ground-truth programs in most networks, and it is the weakest in network 2, whereas the displacement 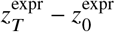 shows no block structure and the KL vector and the displacement 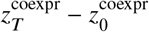 show it only in some networks. All three programs are active at the final timepoint, so an endpoint comparison does not show when each program changes; per-transition KL vectors do (Table 3).

In the switch (Fig. 7), the OT residual and the displacement 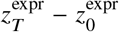 separate the two programs in all six networks, and the KL vector does so in four of them. In networks 1 and 5 neither the KL vector nor the displacement 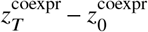 separates the programs. KL captures the coordinated reorganisation of neighbourhoods, and OT captures the accompanying coordinated change in marginal expression as one program is activated and the other suppressed. The switch therefore contains both components of program change rather than pure rewiring.

### Robustness to dropout

Dropout is varied over *p*_*d*_ ∈ [0, 0.5], and the reported quantity is the gap between within-program and between-program cosine similarity of the endpoint vectors, averaged over the six networks of each scenario (Fig. 8). In the switch, both signals separate the programs at low dropout. The OT gap is above that of its displacement 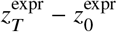 only at low dropout and the two are similar from *p*_*d*_ = 0.2, whereas the KL gap stays above that of the displacement 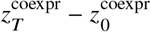 at all dropout levels. Both signals lose about half of their gap between no dropout and *p*_*d*_ = 0.5. In the cascade, the OT gap decreases quickly but stays above that of the other vectors, whereas the KL vector and the displacement 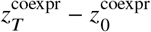 are close to zero from *p*_*d*_ = 0.2. Both signals therefore lose information as dropout increases, and the KL signal does so faster in the cascade and at a similar rate in the switch. The spread between networks is large, so the curves describe the typical trend, not each network. Under strong dropout many of the correlation graphs have no pair above the threshold and the graph is built from the strongest pairs (Appendix A), so these curves describe a regime of very sparse correlation graphs.

**Figure 8:**
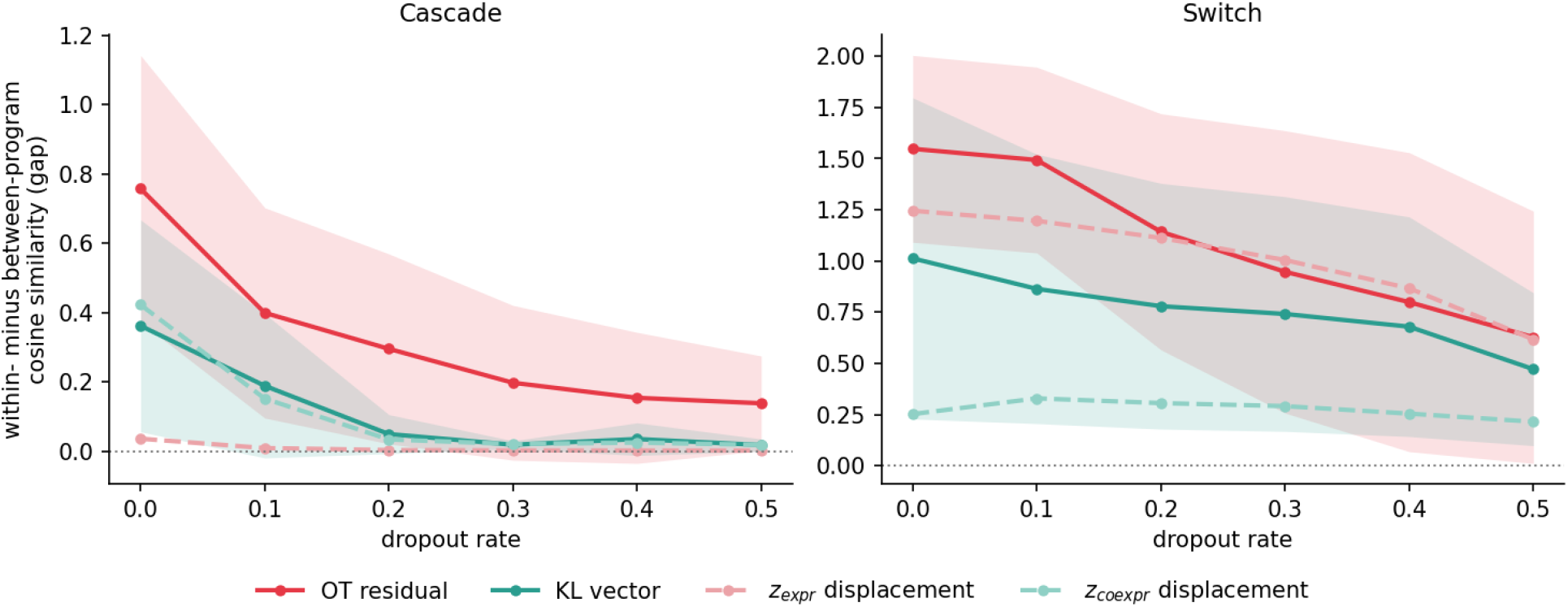
Within-minus between-program cosine similarity of the endpoint vectors versus dropout, in the six cascades (left) and the six switches (right). Solid lines: OT residual and KL vector; dashed lines: displacements 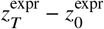 and 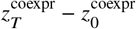. Lines: mean over networks; bands: SD between networks.

**Figure 9:**
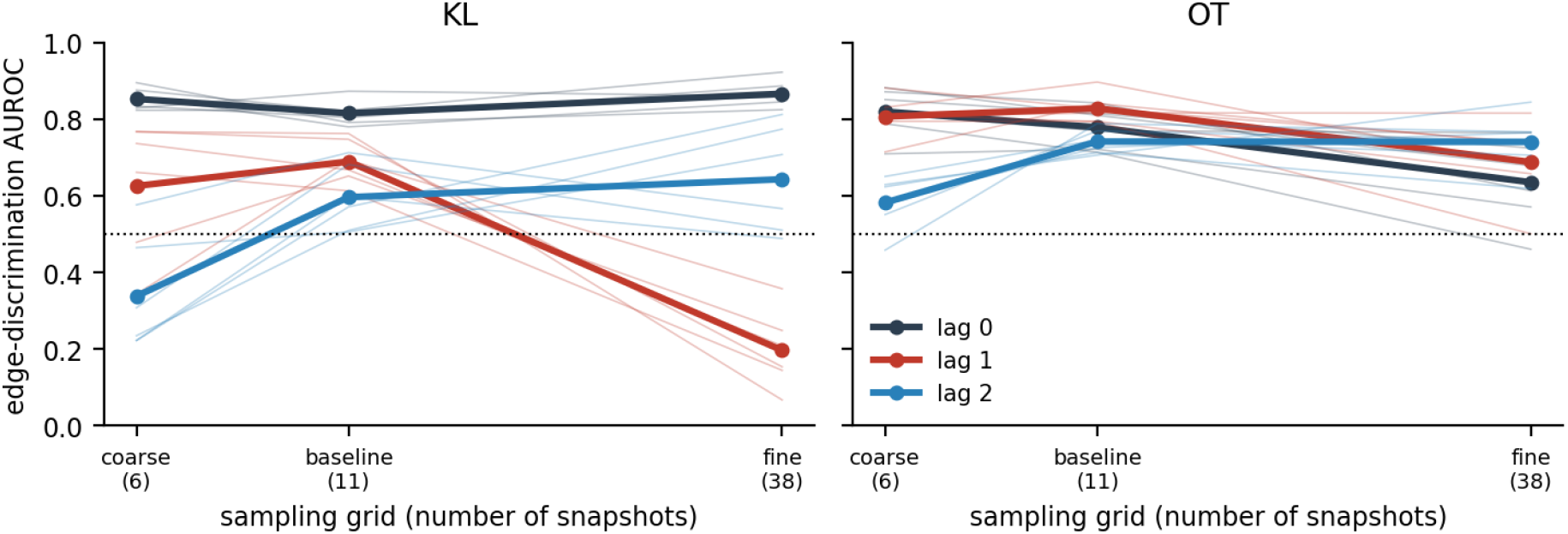
Sensitivity of the lagged channels to the sampling grid. Edge-discrimination AUROC (forward reading, averaged over the two transitions and three seeds) of lags 0, 1 and 2 of KL (left) and OT (right) when the same six cascades are sampled on the coarse (6 snapshots), baseline (11) and fine (38) grid. Thin lines: single networks; thick lines: mean over the six networks. Dotted line: chance. For KL, only lag 0 is informative on every grid; lag 1 falls below chance on the fine grid. Per-network values are in Table 6.

### All representations

Table 3 gives program recovery for all representations on the six cascades and six switches of §3.1. For each representation, the cosine-distance matrix between genes is embedded by multidimensional scaling and clustered by Ward hierarchical clustering at the true number of programs; ARI and NMI compare the clusters with the true programs, and AUROC scores the cosine similarity for same-program against different-program gene pairs. Besides the representations of the main text, the table contains the following variants of the concatenations, which are ablations and not baselines:

- the endpoint OT residual *δ*^OT^ endpoint, computed between the first and the last snapshot (there is no endpoint-only KL variant, since resolving sequential activation requires per-transition vectors, as shown above);
- the full-context variants, which concatenate *z*_concat_ with *δ*^OT^ concat or with *δ*^KL^ concat;
- three max-rank combinations, in which the distance between two genes is the larger of its normalised ranks in two distance matrices: *δ*^OT^ endpoint with *δ*^OT^ concat, *δ*^OT^ concat with *δ*^KL^ concat, and *δ*^OT^ endpoint with *δ*^KL^ concat. Concatenating the *δ* signals with *z*_concat_ does not improve on *z*_concat_ alone. The rank combination of the endpoint OT residual with *δ*^KL^ concat is the only combination that matches the best representation in both scenarios. ARI penalises a few outlying genes that Ward clustering splits off, whereas AUROC reflects all gene pairs, which is why the two can differ.

## D Lagged association: direction and sampling resolution

Table 4 gives the edge discrimination of each channel, Table 5 the direction of the lagged channels, and Table 6 the edge discrimination by sampling resolution and lag; all three are summarised in §3.4. Lagged channels are read forward: the score of a pair is *C*_*i j*_ with *i* in the earlier program and *j* in the later one.

## E CardamomOT on retrieved contexts: all budgets and further baselines

AUPRC is compared with its chance level, the edge base rate. Gene retrieval, the channel ablation and the CardamomOT experiments use a single training seed. DELVE fills the whole budget without program cores and is run at a budget of 30 genes; the lagged-correlation selection is run on the 150-gene cascades.

Table 7 extends Table 2 to all budgets on the 60-gene and 150-gene cascades with TF-to-any relays, with AUPRC/base and, at a budget of 30 genes, the further baselines described after the table. The switches and the cascades with TF-to-TF relays are in Appendix G.

### Further retrieval baselines

In addition to the baselines of the main text, genes are selected with the same procedure as GSET (program cores plus bridges, the same budget) from a single channel: KL at lag 0 alone (Appendix F), the OTVelo gene–gene score [43], and the lagged OT affinity of GeneTrajectory [27, 11] (“OT similarity”). For OTVelo and OT similarity, the score returned by the method replaces *C*_*c*_ in the bridge score of §2.6, so a gene is scored by its best partner in the adjacent program, read forward. DELVE [28], run with its default parameters, uses no programs: it first detects modules of dynamic genes, builds a graph of all cells pooled across timepoints from these genes, and ranks all genes by the Laplacian score on this graph, which is low for genes that vary smoothly along it; its lowest-scoring genes form the context, without cores. These baselines are run at a budget of 30 genes, and their rows are in Table 7. On the 150-gene cascades, DELVE covers no relay edge and the selections from the OTVelo and OT-similarity scores with the programs given cover few, and DELVE selects genes of a single program, like the most variable genes. On the 60-gene cascades, DELVE selects genes of the first two programs and none of the third, so it covers few relay edges of the first transition and none of the second. With the programs given, the OTVelo selection covers few relay edges of the first transition, and the OT-similarity selection covers about half of them, but few when the programs are discovered. Compared with KL at lag 0 alone, the GSET context covers more relay edges of the second transition in every setting, and CardamomOT recovers them at least as well. On the first transition KL alone covers more relay edges on the 60-gene cascades with the programs given and on the 150-gene cascades with the programs discovered.

### Gene-level and network-based selections

GSET is further compared with selections that rank genes individually and with selections from regulatory networks inferred on all genes (Table 8). All of them use the cells of all runs, pooled at each timepoint, and the same budgets as Table 7. *Temporal differential expression* (temporal DE) ranks genes either by the ANOVA F-statistic of their log-expression across timepoints, which compares the variation between timepoints with the variation between cells within a timepoint, or by the variance of their mean log-expression over the timepoints, which favours the genes with the largest change in level. *WGCNA hubs* [19] are the genes of highest module membership in the modules of a WGCNA network built on all cells, with soft-thresholding exponent *β* = 6, the topological overlap matrix and average-linkage clustering cut into as many modules as there are programs, clusters smaller than a minimum size being left unassigned. Each module contributes the same number of hubs, ranked by their correlation with the module eigengene. *Hotspot* [5] ranks genes by the Z-score of their local autocorrelation on a 30-nearest-neighbour graph of the cells built on the first ten principal components of the log-expression, with the depth-adjusted negative binomial model of Hotspot. *GRNBoost2* [24], with all genes or only the TFs as candidate regulators, and *GENIE3* [12], with the TFs as candidate regulators, infer a weighted network on all genes (default parameters of arboreto), and the budget is filled with both endpoints of the edges taken in decreasing order of weight. Two further selections keep the same program cores and bridge budget as GSET. For GRNBoost2, with all genes as candidate regulators, the edge weight *w*_*i j*_ replaces *C*_*c*_ [*i, j*] in the bridge score of §2.6, so a gene is scored by its strongest edge to a gene of the adjacent program. For temporal DE, the bridges are the genes with the highest ANOVA F-statistic, without reference to the programs, and the rest of the budget is filled with the same program cores. GRNBoost2 filled with TF regulators and GENIE3 use the TF annotation, which GSET does not use. On the 150-gene cascades every one of these selections covers fewer relay edges than GSET, with the programs given or discovered. On the 60-gene cascades the variance of the mean expression covers most relay edges of the first transition at a budget of 30 genes, like HVG, and none of the second. GRNBoost2 in the GSET procedure with the programs given covers relay edges of the second transition only, more than GSET on the 60-gene cascades and fewer on the 150-gene cascades. CardamomOT fitted on the contexts of GRNBoost2 in the GSET procedure (Table 7) recovers the relay edges of the first transition less well than on the GSET contexts, at or below chance with the programs given, since these contexts cover none of them. On the second transition it is above GSET on the 60-gene cascades with the programs given and below GSET in the other settings. Over both transitions together these contexts cover fewer relay edges than the GSET contexts in every setting, and on the 60-gene cascades with the programs given the loss on the first transition is much larger than the gain on the second.

**Table 8.** Coverage of relay edges by further gene-level and network-based selections on the cascades with TF-to-any relays (C0→C1 + C1→C2, of six edges each, mean over six networks). The selections are described in the text. Budgets and GSET rows as in Table 7. Rows marked TF use the TF annotation, which GSET does not use.

| Selection | 60 genes |  | 150 genes |  |
|---|---|---|---|---|
|  | budget 18 | budget 30 | budget 30 | budget 45 |
| GSET (true) | 2.3+1.5 | 4.5+3.5 | 5.5+1.3 | 6.0+3.2 |
| GSET (disc.) | 1.0+1.3 | 3.8+3.0 | 2.5+1.5 | 4.2+1.8 |
| Temporal DE, ANOVA | 0.0+0.2 | 0.0+0.3 | 0.0+0.0 | 0.0+0.0 |
| Temporal DE, variance of means | 1.0+0.0 | 4.8+0.0 | 0.0+0.0 | 0.0+0.0 |
| Temporal DE, ANOVA with program cores (true) | 0.0+0.0 | 0.0+0.2 | 0.0+0.0 | 0.0+0.0 |
| Temporal DE, ANOVA with program cores (disc.) | 0.0+0.3 | 0.0+1.7 | 0.0+0.0 | 0.3+0.0 |
| GRNBoost2 fill, all regulators | 0.2+0.0 | 0.7+0.0 | 0.0+0.0 | 0.0+0.2 |
| GRNBoost2 fill, TF regulators | 0.0+0.2 | 1.0+0.2 | 0.0+0.0 | 0.5+0.0 |
| GENIE3 fill, TF regulators | 0.0+0.0 | 0.7+0.2 | 0.8+0.0 | 1.0+0.0 |
| GRNBoost2 in GSET procedure (true) | 0.0+1.8 | 0.0+4.2 | 0.0+1.0 | 0.0+2.2 |
| GRNBoost2 in GSET procedure (disc.) | 0.3+0.7 | 1.0+2.3 | 1.5+0.8 | 2.0+1.2 |
| WGCNA hubs | 0.0+0.0 | 0.0+0.0 | 0.0+0.0 | 0.0+0.0 |
| Hotspot | 0.0+0.0 | 1.2+0.0 | 1.2+0.0 | 1.3+0.7 |

## F Ablation of the retrieval channels

Bridges are selected by KL at lag 0 only, by KL at lag 0 and OT at lag 1 as in GSET (the ratio of the main experiments, with the allocation of §2.6), or by OT at lag 1 only, with the same program cores and the same budget in each family (Table 9; per network on the 150-gene cascades in Table 10). The coverage is compared here. CardamomOT fitted on the KL-only contexts at a budget of 30 genes is in Table 7. Over both transitions, the pair of channels covers more relay edges than KL alone on the cascades with the programs given at budgets of 30 and 45 genes, and on the 60-gene cascades with the programs discovered at a budget of 30 genes. It covers fewer on the 150-gene cascades with the programs discovered and at a budget of 18 genes on the 60-gene cascades, and as many or fewer on the switches. Most of the gain is on the second transition. Without the allocation, a single list ranked by score retrieves mostly sources, since they score higher than targets, and a source covers a relay edge only together with its target. Over three training seeds, the coverage of relay edges at a budget of 30 genes shows the same pattern as with one seed: the pair of channels covers more relay edges than KL alone with the programs given and on the 60-gene cascades with the programs discovered, and fewer on the 150-gene cascades with the programs discovered. Across three encoder seeds (42 to 44), the full GSET context covers 6.8 to 7.5 relay edges (of twelve) on the 150-gene cascades with the programs given and 3.8 to 4.8 with the programs discovered, compared with 1.4 for the best static selection, the TF list, which does not depend on the seed. The advantage over static selection therefore persists across all tested seeds.

**Table 9.**
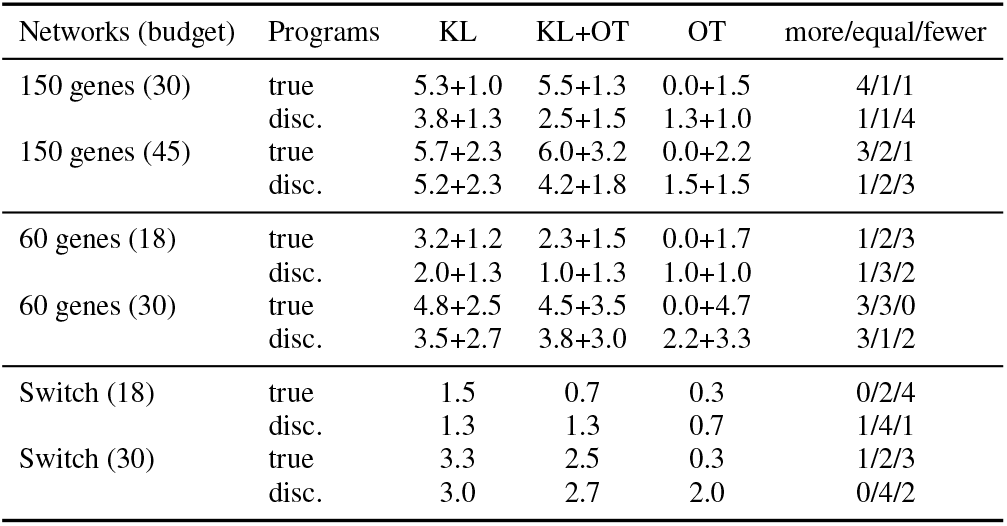
Channel ablation: mean number of covered relay edges over six networks, C0→C1 + C1→C2 for the cascades (of six edges each) and C0→C1 for the switches, with the programs given (true) or discovered (disc.). Last column: number of networks in which KL+OT covers more / as many / fewer relay edges than KL alone (sum over the transitions). Cores and budgets as in Table 7 (150 and 60 genes) and Table 14 (switches).

| Networks (budget) | Programs | KL | KL+OT | OT | more/equal/fewer |
|---|---|---|---|---|---|
| 150 genes (30) | true | 5.3+1.0 | 5.5+1.3 | 0.0+1.5 | 4/1/1 |
|  | disc. | 3.8+1.3 | 2.5+1.5 | 1.3+1.0 | 1/1/4 |
| 150 genes (45) | true | 5.7+2.3 | 6.0+3.2 | 0.0+2.2 | 3/2/1 |
|  | disc. | 5.2+2.3 | 4.2+1.8 | 1.5+1.5 | 1/2/3 |
| 60 genes (18) | true | 3.2+1.2 | 2.3+1.5 | 0.0+1.7 | 1/2/3 |
|  | disc. | 2.0+1.3 | 1.0+1.3 | 1.0+1.0 | 1/3/2 |
| 60 genes (30) | true | 4.8+2.5 | 4.5+3.5 | 0.0+4.7 | 3/3/0 |
|  | disc. | 3.5+2.7 | 3.8+3.0 | 2.2+3.3 | 3/1/2 |
| Switch (18) | true | 1.5 | 0.7 | 0.3 | 0/2/4 |
|  | disc. | 1.3 | 1.3 | 0.7 | 1/4/1 |
| Switch (30) | true | 3.3 | 2.5 | 0.3 | 1/2/3 |
|  | disc. | 3.0 | 2.7 | 2.0 | 0/4/2 |

**Table 10.** Channel ablation on the six 150-gene cascades with TF-to-any relays: number of covered relay edges of the first and the second transition (C0→C1 + C1→C2, of six edges each) for contexts of 30 and 45 genes (10 or 15 program cores; 20 or 30 bridges), with the programs given (upper block) or discovered (lower block). Bridges selected by KL at lag 0 only, by KL at lag 0 and OT at lag 1 as in GSET (14 and 6, or 20 and 10 genes), or by OT at lag 1 only. Rows: networks.

|  | 30 genes |  |  | 45 genes |  |  |
|---|---|---|---|---|---|---|
| Net | KL | KL+OT | OT | KL | KL+OT | OT |
| *Programs given* |  |  |  |  |  |  |
| 1 | 5+2 | 6+2 | 0+0 | 5+4 | 6+3 | 0+0 |
| 2 | 4+4 | 3+4 | 0+3 | 5+4 | 6+6 | 0+4 |
| 3 | 6+0 | 6+1 | 0+0 | 6+3 | 6+2 | 0+3 |
| 4 | 6+0 | 6+1 | 0+1 | 6+0 | 6+3 | 0+1 |
| 5 | 5+0 | 6+0 | 0+3 | 6+1 | 6+3 | 0+3 |
| 6 | 6+0 | 6+0 | 0+2 | 6+2 | 6+2 | 0+2 |
| mean | 5.3+1.0 | 5.5+1.3 | 0.0+1.5 | 5.7+2.3 | 6.0+3.2 | 0.0+2.2 |
| *Programs discovered* |  |  |  |  |  |  |
| 1 | 6+1 | 5+1 | 6+1 | 6+1 | 6+1 | 6+1 |
| 2 | 3+3 | 1+4 | 0+1 | 6+6 | 3+4 | 0+1 |
| 3 | 6+0 | 6+0 | 0+1 | 6+1 | 6+2 | 0+1 |
| 4 | 1+2 | 2+2 | 2+1 | 1+4 | 3+2 | 3+4 |
| 5 | 3+0 | 1+0 | 0+0 | 6+0 | 3+0 | 0+0 |
| 6 | 4+2 | 0+2 | 0+2 | 6+2 | 4+2 | 0+2 |
| mean | 3.8+1.3 | 2.5+1.5 | 1.3+1.0 | 5.2+2.3 | 4.2+1.8 | 1.5+1.5 |

### Learned encoders

To test whether the retrieval depends on the learned encoders, each signal was also computed without training, the KL signal on the rows of the co-expression graph and the OT residual on the 49 quantiles of the standardised expression. The programs are given, and each version takes the cores from its own embedding. On the 150-gene cascades the learned encoders improve each channel alone, but in the full GSET context the coverage is about the same at a budget of 30 genes and higher only at 45. On the 60-gene cascades they improve OT but not KL, and GSET covers more without them (Table 11). The edge discrimination of KL at lag 0 is similar with and without the encoders, that of OT at lag 1 is higher without them on the first transition and lower on the second, and the channels of lag 2 fall below chance on the second transition without them (Table 12). The retrieval therefore does not require the learned encoders on these networks, but they improve the coverage in some settings, slightly at the larger budget on the 150-gene cascades and strongly when the gene panel is large: with 1000 unrelated genes added to these cascades the coverage without them is much lower at budgets of 30 and 45 genes (last block of Table 11, Appendix H).

**Table 11.**
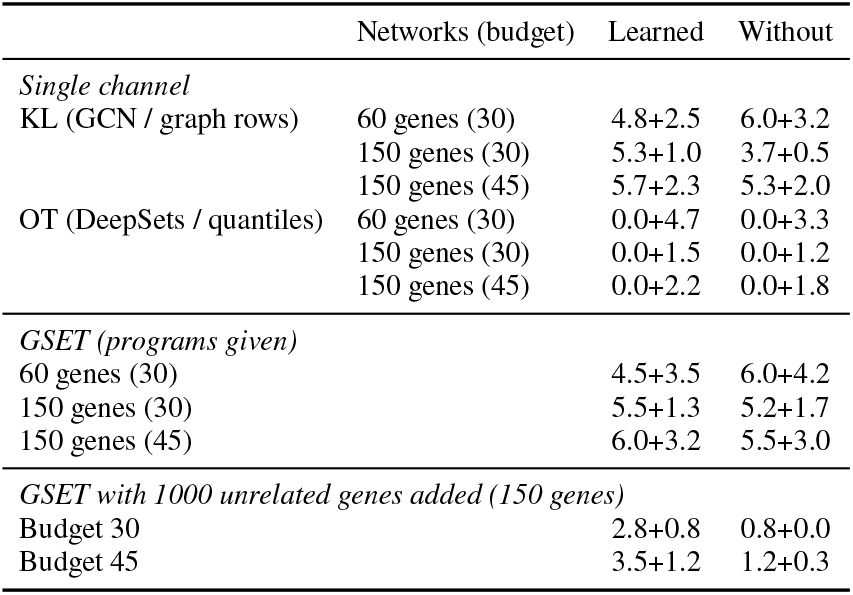
Coverage of relay edges (C0→C1 + C1→C2, of six edges each, mean over six cascades with TF-to-any relays) with the learned encoders of the main experiments (GCN, DeepSets, training seed 42) and without them (rows of the co-expression graph, quantiles of the expression). Five runs of 100 cells per timepoint, budgets and cores as in Table 7. *Single channel*: all bridges come from one channel. *GSET*: KL at lag 0 and OT at lag 1 as in the main experiments. The programs are given, and each version takes the cores from its own embedding. The last block adds 1000 genes of the reprogramming time course to the network (Appendix H), with the same added genes and cells in both versions.

**Table 12.** Discrimination of relay edges on the 60-gene cascades with TF-to-any relays with the learned encoders (as in Table 4, mean over six networks and three training seeds) and without them (KL on the rows of the co-expression graph, OT on the quantiles of the expression, no training). AUROC of true edges against all TF-to-gene pairs without an edge between the same two programs, forward reading. Chance is 0.5.

|  | Learned |  | Without |  |
|---|---|---|---|---|
| Channel | C0→C1 | C1→C2 | C0→C1 | C1→C2 |
| KL, lag 0 | 0.88 | 0.76 | 0.87 | 0.81 |
| KL, lag 1 | 0.78 | 0.60 | 0.89 | 0.68 |
| KL, lag 2 | 0.73 | 0.46 | 0.77 | 0.29 |
| OT, lag 0 | 0.74 | 0.82 | 0.86 | 0.84 |
| OT, lag 1 | 0.81 | 0.85 | 0.88 | 0.78 |
| OT, lag 2 | 0.78 | 0.71 | 0.90 | 0.28 |

## G Benchmarks with TF-to-TF relays (oracle TF list)

Two further families of simulated networks have relay edges that join two TFs, so that the TF list contains both endpoints of every relay edge and acts as an oracle, which makes them easier than the TF-to-any cascades of the main text. The first is a family of five 150-gene cascades with three programs of 50 genes, two TFs per program and four relay edges per transition. GSET covers all relay edges of the first transition with 20% of the genes, while HVG and program cores cover none and random sets almost none, and the TF list, which knows both endpoints, remains above GSET on the second transition (Fig. 10, Table 13). On all edges within the context, the GSET context performs like the full-gene model. The second is the family of switches with edges between the programs (the first program activates the second, which then suppresses the first), with six TFs per program. Its relay edges are the six activating TF-to-TF edges from the first to the second program, and the 36 inhibitory edges back are not evaluated by CardamomOT. Its retrieval is shown in Fig. 3g–i and discussed in §3.6. CardamomOT on its contexts is in Table 14.

**Table 13.**
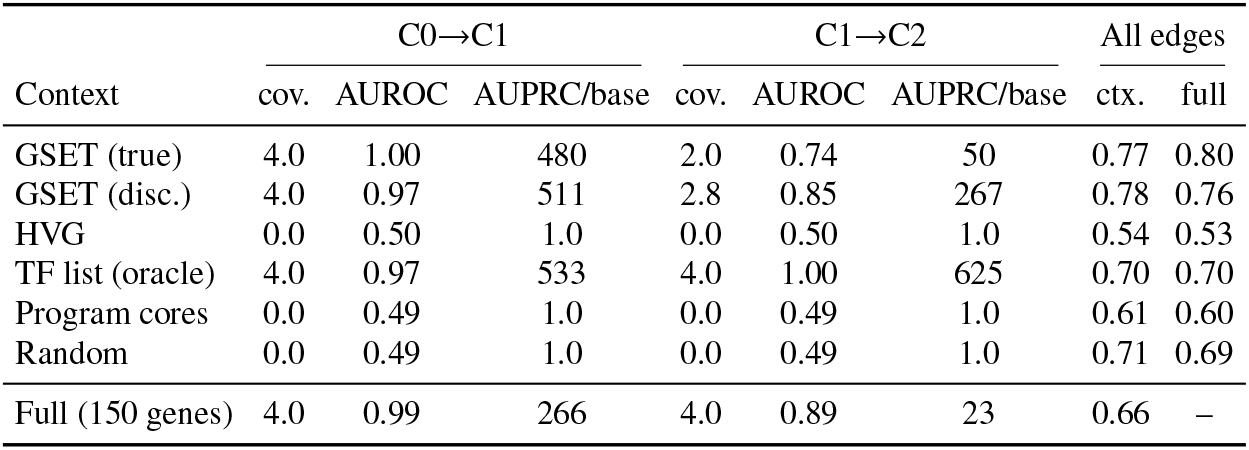
CardamomOT on retrieved contexts on the family of 150-gene cascades with TF-to-TF relays (five networks; three programs of 50 genes, two TFs per program, four TF-to-TF relay edges per transition; relay-edge base rate 0.0016). Budget 30: 10 cores and 20 bridges (14 selected by KL at lag 0 and 6 by OT at lag 1). Columns and rows as in Table 7. The TF list contains both endpoints of every relay edge (oracle).

|  | C0→C1 |  |  | C1→C2 |  |  | All edges |  |
|---|---|---|---|---|---|---|---|---|
| Context | cov. | AUROC | AUPRC/base | cov. | AUROC | AUPRC/base | ctx. | full |
| GSET (true) | 4.0 | 1.00 | 480 | 2.0 | 0.74 | 50 | 0.77 | 0.80 |
| GSET (disc.) | 4.0 | 0.97 | 511 | 2.8 | 0.85 | 267 | 0.78 | 0.76 |
| HVG | 0.0 | 0.50 | 1.0 | 0.0 | 0.50 | 1.0 | 0.54 | 0.53 |
| TF list (oracle) | 4.0 | 0.97 | 533 | 4.0 | 1.00 | 625 | 0.70 | 0.70 |
| Program cores | 0.0 | 0.49 | 1.0 | 0.0 | 0.49 | 1.0 | 0.61 | 0.60 |
| Random | 0.0 | 0.49 | 1.0 | 0.0 | 0.49 | 1.0 | 0.71 | 0.69 |
| Full (150 genes) | 4.0 | 0.99 | 266 | 4.0 | 0.89 | 23 | 0.66 | – |

**Table 14.** CardamomOT on retrieved contexts on the switches (six networks, 60 genes; two programs of 30 genes, six TFs each). The relay edges are the six activating TF-to-TF edges from the first to the second program (relay-edge base rate 0.0067); the inhibitory edges back are not evaluated. All edges join TFs, so the TF list is an oracle. Budgets 18 and 30 use 6 cores and 12 or 24 bridges. Columns and rows as in Table 7.

|  | Budget 18 |  |  |  |  | Budget 30 |  |  |  |  |
|---|---|---|---|---|---|---|---|---|---|---|
|  | C0→C1 |  |  | All edges |  | C0→C1 |  |  | All edges |  |
| Context | cov. | AUROC | AUPRC/b | ctx. | full | cov. | AUROC | AUPRC/b | ctx. | full |
| GSET (true) | 0.7 | 0.51 | 3.2 | 0.64 | 0.63 | 2.5 | 0.60 | 4.0 | 0.61 | 0.61 |
| GSET (disc.) | 1.3 | 0.58 | 26 | 0.75 | 0.67 | 2.7 | 0.62 | 13.5 | 0.68 | 0.68 |
| HVG | 0.0 | 0.50 | 1.0 | 0.53 | 0.51 | 0.0 | 0.46 | 1.0 | 0.50 | 0.53 |
| TF list (oracle) | 6.0 | 0.95 | 125 | 0.63 | 0.60 | 6.0 | 0.99 | 125 | 0.63 | 0.60 |
| Program cores | 0.0 | 0.48 | 1.0 | 0.57 | 0.46 | 0.0 | 0.43 | 1.0 | 0.54 | 0.47 |
| Random | 0.2 | 0.49 | 1.2 | 0.66 | 0.61 | 0.7 | 0.47 | 3.7 | 0.67 | 0.64 |
| Lag-corr. (true) | 0.0 | 0.48 | 1.0 | 0.65 | 0.67 | 0.0 | 0.43 | 1.0 | 0.60 | 0.61 |
| Lag-corr. (disc.) | 0.8 | 0.54 | 9.8 | 0.72 | 0.70 | 1.8 | 0.58 | 23.6 | 0.71 | 0.69 |
| Full (60 genes) | 6.0 | 0.63 | 7.9 | 0.64 | – | 6.0 | 0.63 | 7.9 | 0.64 | – |

**Figure 10:**
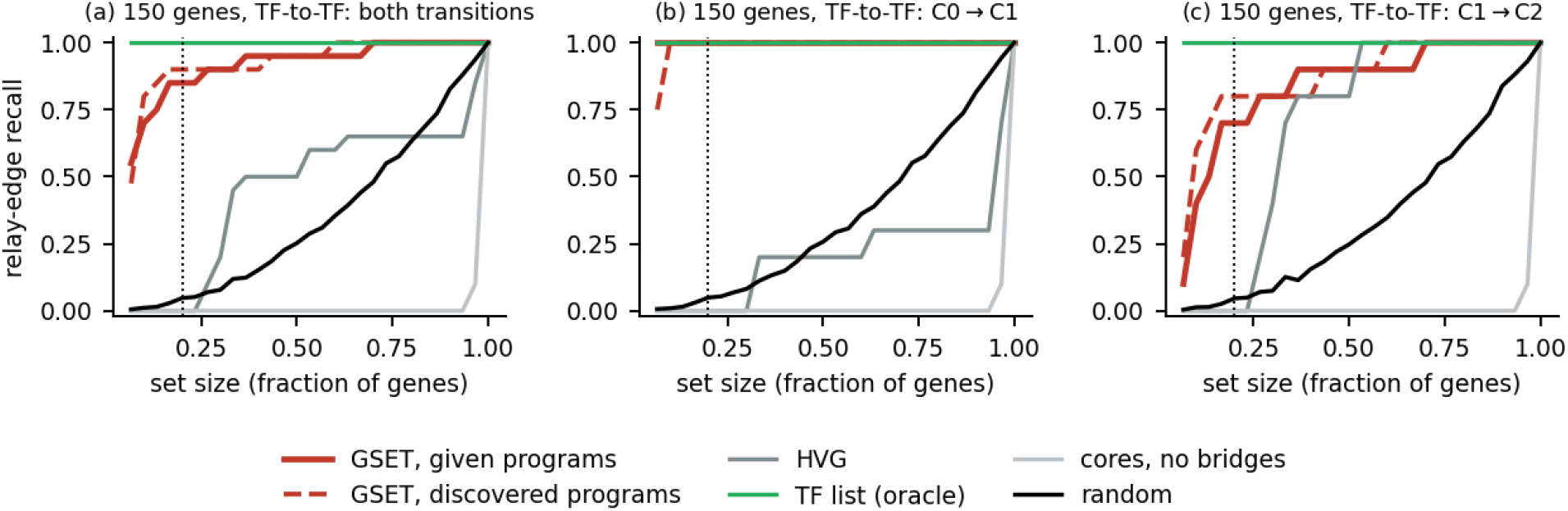
Retrieval of bridges on the 150-gene cascades with TF-to-TF relays (five networks): relay-edge recall as a function of the size of the selected set, for both transitions together (a), C0→C1 (b) and C1→C2 (c). GSET bridges (KL at lag 0 and OT at lag 1 in a ratio of about 70 to 30, without cores), with the programs given (solid) and discovered (dashed), are compared with HVG, the TF list, program cores without bridges and random subsets of the same size. The TF list contains both endpoints of every relay edge (oracle). The dotted line marks the budget used with CardamomOT in Table 13.

## H Robustness to unrelated genes

To test how the retrieval depends on the size of the gene panel, 1000 real genes unrelated to the network were added to each of the six 150-gene cascades with TF-to-any relays. The added genes come from the reprogramming time course of Schiebinger et al. [31] (Data (iii)). For every run and timepoint of the simulation, as many cells as simulated cells were drawn from this data set (days 0–3, matched to the timepoints of the simulation by rank), and the log-normalised expression of 1000 genes chosen at random among its 4,007 genes was appended to the columns of the simulated cells, with the same genes for all matrices. Since the simulated and real cells are sampled independently, the added genes have no designed within-cell correlations with the network genes, although weak correlations may arise by chance in finite samples. They belong to none of the programs. The programs of the network genes are given, the encoders are trained on all 1150 genes (training seed 42), and the measure is the number of relay edges covered at the budgets of the main experiments (Table 15).

**Table 15.** Coverage of relay edges (C0→C1 + C1→C2, of six edges each, mean over six 150-gene cascades with TF-to-any relays). Learned encoders, training seed 42, programs given, budgets and cores as in Table 7. *1000 added genes*: unrelated genes of the reprogramming time course added to the network. *GSET (MR)*: the same method with the cosine similarity of KL at lag 0 replaced by its mutual rank (exploratory). *None*: no genes added. Random: mean of three sets.

|  | 1000 added genes |  |  |  |  |  | None |  |
|---|---|---|---|---|---|---|---|---|
| Budget | GSET | GSET (MR) | HVG | TF list | Cores | Random | GSET | GSET (MR) |
| 30 | 2.8+0.8 | 3.8+1.5 | 0.0+0.0 | 0.7+0.7 | 0.0+0.0 | 0.0+0.0 | 5.5+1.3 | 5.3+1.7 |
| 45 | 3.5+1.2 | 4.7+2.2 | 0.0+0.0 | 0.7+0.7 | 0.0+0.0 | 0.0+0.0 | 6.0+3.2 | 6.0+3.3 |

GSET covers fewer relay edges than without the added genes, but still clearly more than the static selections, which cover at most 0.7 relay edges per transition. The budget is the same as without the added genes, so it is now a much smaller share of the panel. Although the added genes do not belong to the simulated programs, their inclusion changes the embeddings and the neighbourhood structure and introduces additional candidates during bridge retrieval. The reduction in coverage therefore reflects the sensitivity of the retrieval to the expanded gene panel. One possible source is hubness, in which a few genes are similar to many others and take the top ranks of the cosine similarity. As an exploratory variant, introduced after this reduction was observed and in the spirit of hubness-reduction methods, the cosine similarity of KL at lag 0 was replaced by its mutual rank 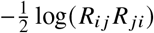, with the embeddings, the OT channel, the cores and the budget unchanged, where *R*_*i j*_ is the rank of gene *j* among the neighbours of gene *i*. With the added genes this variant covers more relay edges, and without them it covers about as many as the cosine (Table 15). Without the learned encoders the coverage with the added genes is much lower (Table 11), so the encoders matter when the gene panel is large.

## I HARISSA simulations: design and aggregation over runs

Each network is simulated with 100 cells per timepoint in five independent runs (three for the switch and for the cascades with TF-to-TF relays), sampled at eleven timepoints between 0 and 300 in simulation time units (ten between 0 and 240 for the switch). The coarse and fine grids of §3.4 sample the same time course at six and thirty-eight timepoints, at constant steps of 60 and 8 time units. The encoders are trained on the matrices of all runs.

The 60-gene scenarios have six independently generated networks each. In the cascade, three programs of 20 genes, each with six TFs, are activated sequentially by a stimulus, and six relay edges per transition join the TFs of one program to any gene of the next; the activation order is verified for each network. In the switch, two programs of 30 genes are simulated in two phases, with the stimulus acting on C0 and then on C1. The 150-gene scenario consists of six cascades of the same design, with programs of 50 genes and five TFs per program.

Both 60-gene scenarios are used for program recovery (§3.3), the 60-gene cascade for the association analysis and the resolution sweep (§3.4), and the 60-gene and 150-gene cascades for retrieval with CardamomOT (§3.6).

### Aggregation over runs

For program recovery, the cells of all runs are pooled at each timepoint; the endpoint maps of Appendix C are computed on a single run. For the association analysis and the retrieval, the per-transition signal vectors are computed separately for each run and averaged over the runs before the lagged similarities are computed. The embeddings in *z*_concat_ and the scores of OTVelo and OT similarity are averaged over the runs in the same way, and the lagged-correlation baseline uses mean profiles averaged over the runs. CardamomOT, HVG, the TF list and DELVE use the cells of all runs.

### Single run

With the signals of a single run of 100 cells per timepoint instead of the average over five runs, GSET covers slightly fewer relay edges on the 150-gene cascades with the programs given (5.8 instead of 6.8 of twelve at a budget of 30 genes, 7.8 instead of 9.2 at 45) and as many or more with the programs discovered, still far more than the TF list, which covers the same 1.4 relay edges on a single run.

## J Scalability

Wall-clock time and memory are measured for GSET and the comparison methods along two axes, the number of genes (RMS, 13,968 cells) and the number of cells (the full Schiebinger time course, 37 time points). Genes are the top-variance genes, chosen deterministically, so every row uses a nested, reproducible gene set. All runs use one machine (10 CPU cores, 34 GB RAM) with nothing else running; a method is timed from the loaded data to its final output. GSET is split into its two parts: GSET_KL_ (the graph network on the co-expression graph, which includes the dense *n* × *n* gene graph) and GSET_OT_ (the DeepSets encoder, linear in cells and genes). Memory is the increase over the process that has just loaded the data.

### Genes

GSET scales mildly with the number of genes (Table 16). The KL part is almost constant (about 6 s up to 1,000 genes) until the dense graph dominates at all 8,442 genes (878 s), and the OT part grows roughly linearly (23 s at 50 genes, 485 s at 1,000), so the total is 492 s at 1,000 genes and 98 minutes at all 8,442 genes, using 1.7 GB (1,000 genes) and 10.2 GB (8,442 genes) above baseline. CardamomOT on the full gene set grows much faster: 951 s at 50 genes, 9,680 s at 200 and 35,595 s (9.9 h) at 400 genes, about 90 and 160 times the time of GSET at 200 and 400 genes; it did not complete at 1,000 genes (see the caption). OT similarity grows roughly quadratically in the number of genes, since it solves one transport problem per gene pair, reaching 2,249 s at 400 genes, and has no valid run at 1,000 genes. OTVelo has an almost constant run time (about 40 s up to 1,000 genes), but its memory grows with the number of genes.

**Table 16.** Gene axis on RMS (13,968 cells), top-variance genes. Time / extra memory above the loaded-data baseline. CardamomOT is run on the full gene set (all genes, with degradation rates); its memory was not recorded in this series. OT similarity at 1,000 genes: the Sinkhorn kernel underflows, all transport solves return zero, and the reported run time of 63 s is therefore not valid (a valid run is estimated at about 9 h by extrapolation). CardamomOT did not complete at 1,000 genes on this machine (the process was killed by the system while memory and swap were nearly exhausted). OT similarity at 8,442 genes did not complete.

| Genes | GSET<sub>KL</sub> | GSET<sub>OT</sub> | GSET total | OTVelo | OT similarity | CardamomOT |
|---|---|---|---|---|---|---|
| 50 | 6 s | 23 s | 29 s / 0.3 GB | 43 s / 1.2 GB | 69 s / 0.5 GB | 951 s |
| 100 | 6 s | 43 s | 49 s / 0.2 GB | 40 s / 1.2 GB | 273 s / 0.5 GB | 3,902 s |
| 150 | 6 s | 80 s | 86 s / 0.3 GB | 40 s / 1.3 GB | 477 s / 0.5 GB | 7,020 s |
| 200 | 6 s | 101 s | 107 s / 0.3 GB | 38 s / 1.3 GB | 601 s / 0.6 GB | 9,680 s |
| 400 | 6 s | 222 s | 228 s / 0.8 GB | 43 s / 1.5 GB | 2,249 s / 0.7 GB | 35,595 s |
| 1,000 | 7 s | 485 s | 492 s / 1.7 GB | 39 s / 2.1 GB | numerical failure | not completed |
| 8,442 | 878 s | 5,023 s | 5,902 s / 10.2 GB | 53 s / 11.3 GB | not completed | n/a |

### Cells

At 50 genes all methods can be compared (Table 17). GSET is linear in the number of cells: the OT part grows from 31 s to 93 s over a fourfold increase in cells and the KL part does not depend on the cells (33 s, since it works on the gene graph), so the total grows from 65 s to 126 s. CardamomOT grows faster than linearly in the number of cells (1,715 s, 3,222 s and 8,334 s), from 26 to 66 times the time of GSET. OT similarity is almost independent of the number of cells (186 to 206 s). OTVelo grows faster than linearly (8 s to 151 s from 25% to 100% of the cells). At 1,000 genes, where only GSET and OTVelo run (Table 18), GSET is again linear (646 s, 1,078 s, 2,021 s) and its memory increase stays moderate (1 to 3 GB).

**Table 17.** Cell axis on Schiebinger (37 time points), 50 genes, random subsets of cells. Time / extra memory above the loaded-data baseline. CardamomOT is run on the 50 genes as its full context. Its memory is noisy and does not grow steadily with the number of cells.

| Cells | GSET<sub>KL</sub> | GSET<sub>OT</sub> | GSET total | OTVelo | OT similarity | CardamomOT |
|---|---|---|---|---|---|---|
| 12,724 (25%) | 34 s | 31 s | 65 s / 0.4 GB | 8 s / 0.2 GB | 206 s / 0.5 GB | 1,715 s / 2.5 GB |
| 25,448 (50%) | 33 s | 48 s | 81 s / 0.5 GB | 31 s / 1.5 GB | 186 s / 0.5 GB | 3,222 s / 1.9 GB |
| 50,897 (100%) | 33 s | 93 s | 126 s / 0.5 GB | 151 s / 1.5 GB | 203 s / 0.6 GB | 8,334 s / 3.7 GB |

**Table 18.**
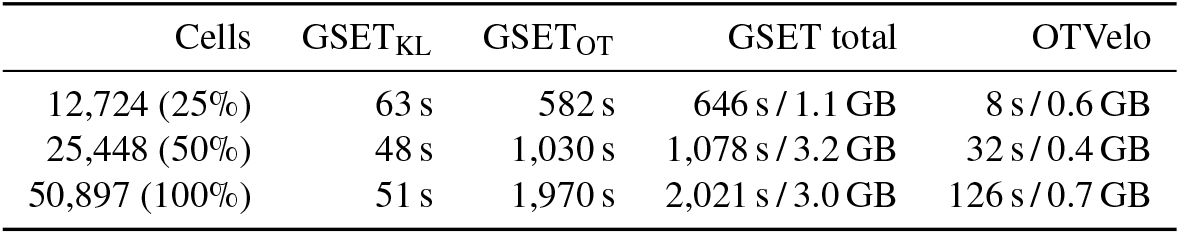
Cell axis on Schiebinger (37 time points), 1,000 genes, random subsets of cells. Time / extra memory above the loaded-data baseline. Memory increases are approximate and not strictly monotone. OT similarity did not give a valid result already at 25% of the cells and was not run on larger subsets.

| Cells | GSET<sub>KL</sub> | GSET<sub>OT</sub> | GSET total | OTVelo |
|---|---|---|---|---|
| 12,724 (25%) | 63 s | 582 s | 646 s / 1.1 GB | 8 s / 0.6 GB |
| 25,448 (50%) | 48 s | 1,030 s | 1,078 s / 3.2 GB | 32 s / 0.4 GB |
| 50,897 (100%) | 51 s | 1,970 s | 2,021 s / 3.0 GB | 126 s / 0.7 GB |

**Table 19.** Lagged retrieval on days 0–3 of the reprogramming time course for three training seeds. The score of the query gene and its 29 top retrieved genes is computed in every cell. *Day*: median day of the top 10% of cells by score. *AUROC*: separation of dividing cells (top quarter of the cell-cycle signature of Schiebinger et al. [31]) from the other cells by the score. *S/G2M*: number of S and G2/M genes [34] among the top 30. Bold: see Evaluation above.

|  |  | Seed 42 |  |  | Seed 43 |  |  | Seed 44 |  |  |
|---|---|---|---|---|---|---|---|---|---|---|
| Query | Channel | Day | AUROC | S/G2M | Day | AUROC | S/G2M | Day | AUROC | S/G2M |
| Lox | KL, lag 1 | **2.5** | **0.66** | **5** | 2.5 | 0.63 | 4 | **2** | **0.82** | **7** |
| Bgn | KL, lag 1 | 2.5 | 0.37 | 0 | **2.5** | **0.66** | **7** | **2** | **0.73** | **6** |
| Cyr61 | KL, lag 1 | **2.5** | **0.67** | **5** | **2.5** | **0.69** | **7** | 1.5 | 0.64 | 2 |
| Col1a1 | KL, lag 1 | 2.5 | 0.36 | 0 | 2.5 | 0.57 | 4 | 2 | 0.73 | 2 |
| Lox | OT, lag 2 | 2 | 0.75 | 2 | 2 | 0.81 | 4 | 2 | 0.82 | 3 |
| Bgn | OT, lag 2 | **1.5** | **0.82** | **7** | **2** | **0.85** | **9** | **2** | **0.87** | **8** |
| Cyr61 | OT, lag 2 | 1 | 0.53 | 1 | 0.5 | 0.38 | 0 | 1 | 0.42 | 0 |
| Col1a1 | OT, lag 2 | **1.5** | **0.81** | **6** | **2** | **0.86** | **9** | **2** | **0.85** | **9** |

### CardamomOT on retrieved contexts

CardamomOT is the most expensive method, and its cost depends on the number of genes it is given, not on how the genes were chosen. On the 50-gene context retrieved by GSET for HSPA1B, CardamomOT takes 851 s (14 minutes), independent of the size of the full transcriptome. With the retrieval time added, GSET followed by CardamomOT on the retrieved context takes about 1,340 s at 1,000 genes and about 6,750 s (1.9 h) at all 8,442 genes, whereas CardamomOT on the full gene set needs 35,595 s at 400 genes and did not complete at 1,000 genes. Retrieval therefore makes the mechanistic model feasible at transcriptome scale instead of replacing it.

## K Reprogramming time course: data and seeds

### Data

The serum condition of Schiebinger et al. [31] (GSE122662) is restricted to days 0–3 (seven timepoints at half-day steps, 34,101 cells). After removing mitochondrial, ribosomal and rarely expressed genes, the 4,000 most variable genes are used, together with seven genes kept for evaluation (4,007 genes). The encoders are trained on this window only.

### Evaluation

The query genes are extracellular matrix genes. Ten genes were tested (Col1a1, Serpine1, Ctgf, Cyr61, Inhba, Ccl2, Thbs1, Lox, Timp3, Bgn); the four shown are those that pass the criterion below in at least one channel and seed, and each channel passes it in 6 of the 30 combinations of gene and seed. All ten are expressed more highly at day 0 than in the later cells (AUROC of the expression for the cells of day 0 between 0.86 and 0.98; 0.89 to 0.94 for the four shown). Lox and Bgn are not in the gene list of the MEF-identity signature of Schiebinger et al. [31], but their expression is among the most correlated with this signature across the cells of days 0–3 (second and fourth of 4,007 genes). The query gene and its 29 top retrieved genes are scored in every cell with Scanpy score genes. Dividing cells (cells with a high cell-cycle signature in Fig. 2) are the top quarter of cells for the cell-cycle signature of Schiebinger et al. [31]. Table 19 marks in bold the retrievals in which the median day of the top 10% of cells by score lies between 1 and 2.5, their mean cell-cycle signature is above the mean of the window, and the top 30 genes include at least five S or G2/M genes of Tirosh et al. [34].

### Results

Without lag, KL retrieval returns the day-0 program in all seeds. At lag 1, KL meets the criterion for Lox, Bgn and Cyr61 in two of three seeds each (bold in Table 19), but not for Col1a1. Its top genes never overlap with those retrieved without lag. At lag 2, OT meets the criterion for Bgn and Col1a1 in all three seeds, with S-phase genes such as Mcm4, Mcm5 and Mcm6 and G2/M genes such as Ckap2l and Ncapd2 among the top 30. From Lox it reaches the dividing cells but retrieves fewer S and G2/M genes, and from Cyr61 it does not reach them. The two channels thus reach the cell-cycle program from partly different genes (Table 19). The mean-expression baseline does not reach dividing cells and retrieves no S or G2/M genes. OTVelo [43], run on the same window with its lag of one transition and its gene–gene score summed over transitions and read forward from the query gene, retrieves the extracellular matrix genes of day 0 from both Lox and Bgn and does not reach dividing cells. Under the weaker criterion that the score separates dividing cells from the others (AUROC above 0.5), KL at lag 1 does so in at least two of three seeds from eight of the ten genes and OT at lag 2 in all seeds from three (Col1a1, Lox and Bgn).

### Bridge retrieval

The fibroblast and cell-cycle programs are those of Lox and of Birc5, an S/G2M gene that Lox reaches by lagged KL in all seeds, each the gene and the 29 genes most similar to it in *z*_concat_ (the two programs can share genes, and the bridges are taken outside their union). The other genes are assigned to the nearer program by centroid in *z*_concat_, and 200 bridges (140 by KL at lag 0, 60 by OT at lag 1, the default channels fixed in advance as in §2.6 and not chosen after seeing these data, without cores) are selected outside the two programs. Pairs of training seeds share 54, 51 and 45 of the 200 genes (about 10 expected by chance). The bridges include subunits of the SAGA (TADA1, TADA3, SGF29, TAF6; recurring in two of three seeds) and NuA4/TIP60 (MRGBP, ING3, MEAF6, DMAP1) acetyltransferase complexes; Gcn5, the catalytic subunit of SAGA, is required for the initiation of reprogramming [9]. Histone acetyltransferase terms are the top terms of an enrichment analysis (GO, Reactome, KEGG, Hallmark) in all three seeds, but their enrichment is not significant after correction for multiple testing (*q* = 0.07 to 0.18). The bridges are candidates without ground truth.

## L RMS experimental-data validation: extended detail

On the RMS data (§3.1), lagged KL is tested by whether increasing the lag from a quiescence-associated query gene traces the known quiescent→intermediate→proliferative progression. The six time bins are pseudotime bins, built partly from proliferation markers (see below), and the retrieval does not use the cell-state annotation. HSPA1B is used as the query gene because its expression is concentrated in quiescent cells, which dominate the early timepoints.

Increasing the lag traces the progression for HSPA1B (Fig. 11), with lag-2 retrieval dominated by replication and mitotic machinery (GINS4, MCM10, KIF11, NDC80, SPC24/25, CEP55, BRCA1), in 6 of 7 training seeds (Fig. 12). A baseline on the same bins, which ranks genes by the lagged similarity of their mean-expression profiles, does not reach the proliferative state (Table 21). Independent single-cell studies of RMS tumours and xenografts report that quiescent progenitor-like cells regenerate the proliferative population after chemotherapy [4], which is consistent with the direction of this progression, although in a different setting and without resolving the intermediate state. The progenitor marker CD44 [4] reproduces the step to the intermediate state in six of seven seeds and reaches the proliferative state at lag 2 in three (Fig. 11, Fig. 14, Table 20).

**Table 20.** Query genes on the RMS data. Expr.: AUROC of the expression of the gene for quiescent cells. Number of training seeds (of seven) in which the best-matching state is quiescent at the static level, intermediate at lag 1, proliferative at lag 2, and in which all three hold. Upper block: genes selected by the expression criterion; lower block: progenitor markers of Danielli et al.

| Query | Expr. | static | lag 1 | lag 2 | all |
|---|---|---|---|---|---|
| HSPA1B | 0.80 | 7 | 7 | 6 | 6 |
| EIF1 | 0.87 | 7 | 6 | 3 | 2 |
| CRYAB | 0.85 | 7 | 6 | 2 | 1 |
| NACA | 0.85 | 7 | 5 | 0 | 0 |
| DNAJB1 | 0.82 | 7 | 7 | 5 | 5 |
| TCEAL9 | 0.79 | 7 | 7 | 4 | 4 |
| GAPDH | 0.78 | 7 | 4 | 0 | 0 |
| NPM1 | 0.75 | 7 | 3 | 0 | 0 |
| EEF1B2 | 0.73 | 7 | 3 | 0 | 0 |
| VIM | 0.71 | 7 | 1 | 3 | 0 |
| HSPH1 | 0.70 | 7 | 7 | 7 | 7 |
| CD44 | 0.65 | 7 | 6 | 3 | 2 |
| COL3A1 | 0.56 | 6 | 3 | 3 | 0 |
| FN1 | 0.39 | 0 | 2 | 2 | 0 |
| COL1A1 | 0.26 | 0 | 0 | 3 | 0 |
| EGFR | 0.41 | 0 | 0 | 0 | 0 |
| THY1 | 0.44 | 0 | 3 | 1 | 0 |

**Table 21.** Program-score AUROC of the mean-expression baseline and of the OT channel (seed 42). Fwd: forward reading of the lagged similarity.

|  | Quiescent | Intermediate | Proliferative |
|---|---|---|---|
| *HSPA1B* |  |  |  |
| Mean-expression baseline, lag 0 | **0.91** | 0.27 | 0.33 |
| Mean-expression baseline, lag 1 | **0.82** | 0.40 | 0.27 |
| Mean-expression baseline, lag 2 | **0.58** | 0.46 | 0.46 |
| OT, lag 0 | 0.64 | 0.09 | **0.81** |
| OT, lag 1 fwd | 0.55 | 0.18 | **0.80** |
| OT, lag 2 fwd | **0.80** | 0.37 | 0.32 |
| *MKI67* |  |  |  |
| Mean-expression baseline, lag 0 | 0.44 | 0.16 | **0.93** |
| Mean-expression baseline, lag 1 | **0.70** | 0.15 | 0.67 |
| Mean-expression baseline, lag 2 | **0.89** | 0.33 | 0.29 |
| OT, lag 0 | 0.44 | 0.16 | **0.94** |
| OT, lag 1 fwd | 0.58 | 0.25 | **0.69** |
| OT, lag 2 fwd | **0.58** | 0.54 | 0.36 |

**Figure 11:**
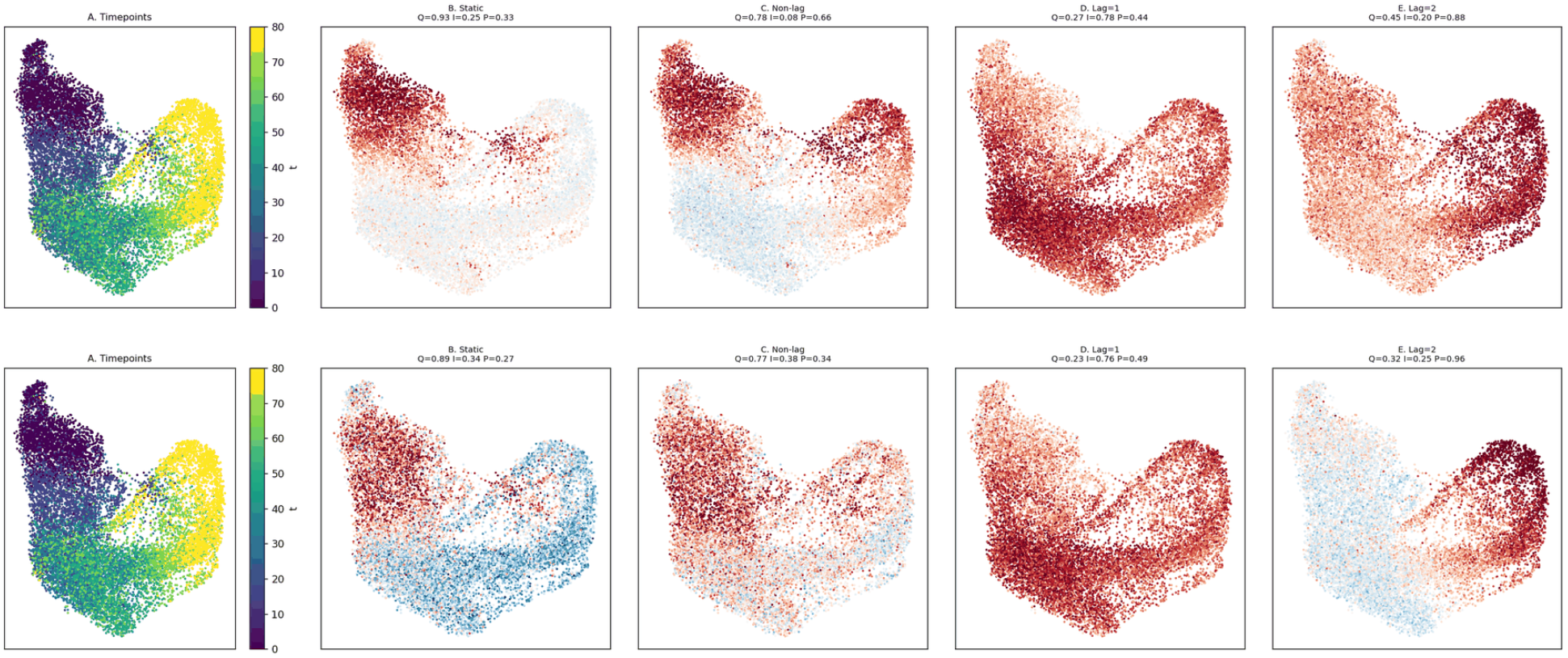
Query genes on RMS data, training seed 42. Top row: HSPA1B; bottom row: CD44, a progenitor marker [4]. **A**: cells coloured by sampling time. **B**: static program, best matching the quiescent state. **C**: same-interval *δ*_KL_ retrieval, quiescent. **D**: lag 1, intermediate. **E**: lag 2, proliferative. Scores are Scanpy score genes on normalised log-expression, with no cell-state annotation used for retrieval. The AUROC against the annotated cell states (Q, I, P: quiescent, intermediate, proliferative) is printed above each panel. Results across training seeds and the MKI67 query are below.

**Figure 12:**
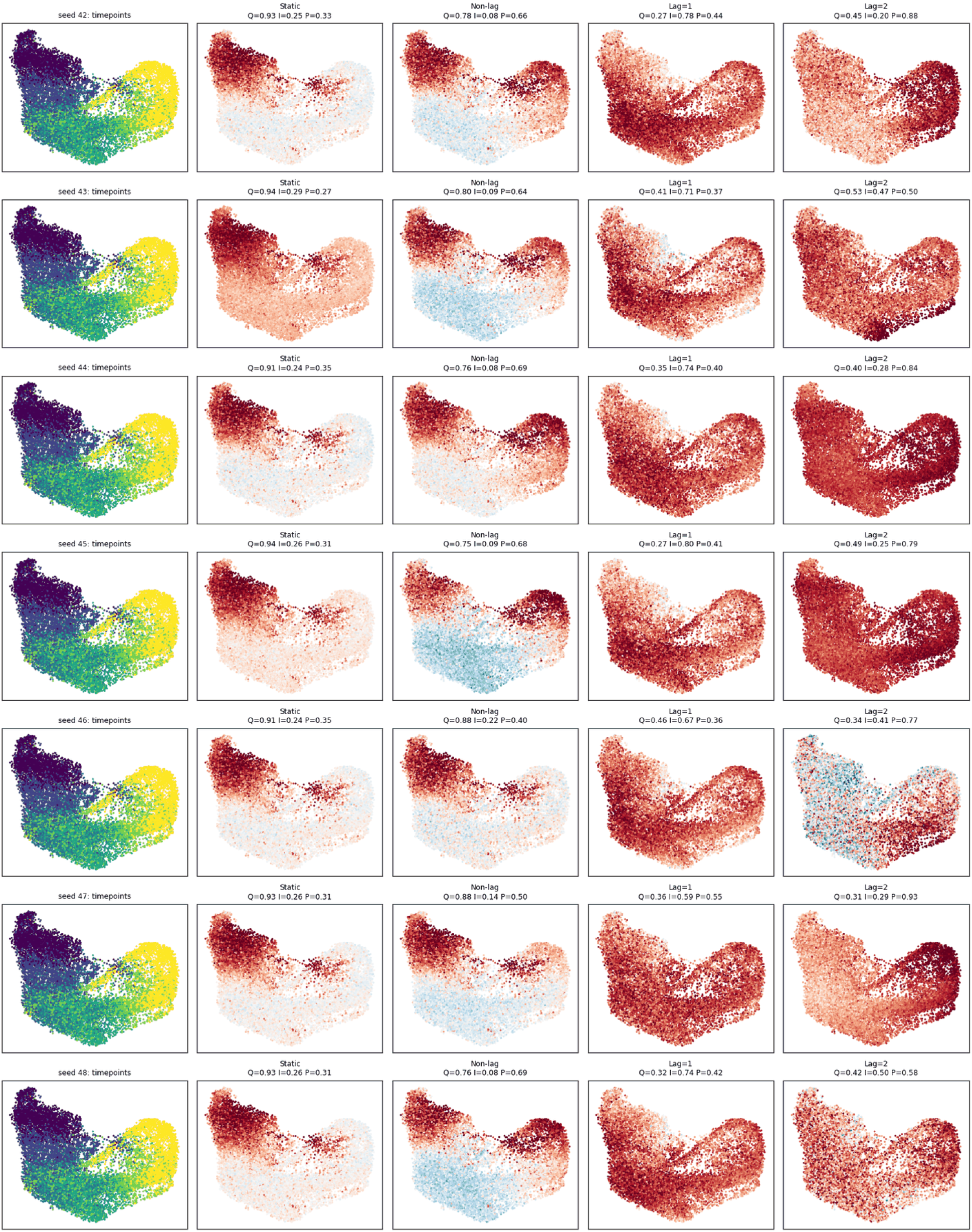
HSPA1B query across 7 GCN training seeds (42–48). Each row shows Timepoints, Static, Non-lag, Lag=1 and Lag=2 program scores for one seed; AUROC (Q/I/P) is reported per panel. The quiescent→intermediate→proliferative shift from Static/Non-lag to Lag=2 is visible in 6 of 7 seeds, in one of them marginally.

### Pseudotime construction

The six timepoints are quantile bins of a pseudotime, not experimental times. Cells of two RMS samples were filtered by median absolute deviation of gene counts, total counts and mitochondrial fraction, normalised, and clustered (Leiden) on the PCA space, and a few clusters were removed. Palantir pseudotime was computed on the PCA space from a start cell of one Leiden cluster (the cell with the largest mean distance to all other cells), normalised to [0, 1] and averaged with equal weight with a normalised proliferation signature (mean *z*-score of *MKI67* and *BIRC5*); the combined score was cut into six quantile bins assigned to *t* = 0, 16, …, 80. The cell-state annotation (quiescent, intermediate, proliferative) is derived separately from marker signatures of the Leiden clusters and is not used by GSET. It is not independent of the pseudotime, however: the start cluster is one of the clusters annotated as quiescent, and the proliferation signature contributes to the ordering. The progression from quiescent to proliferative cells along these bins is therefore partly built into the time axis, and the result should be read as evidence that lagged KL retrieval follows this ordering, not as an independent test of it.

### Scoring

All scores use Scanpy score genes on the library-size-normalised, log-transformed expression matrix, applied to the top 30 genes of each retrieval. Each score is evaluated against the annotated cell states by AUROC (Q, I, P: quiescent, intermediate, proliferative). Four levels of the KL channel are compared for a query gene: static (GCN co-expression similarity), non-lagged co-change (*δ*_KL concat_), and lagged association at lag 1 and lag 2 (forward reading).

### HSPA1B across training seeds

Figure 12 shows the four levels for seven independently trained GCN seeds. The static and non-lagged retrievals are aligned with the quiescent state in all seeds. At lag 1 the intermediate state is the best-matching state in all seven seeds. At lag 2 the proliferative state is the best-matching state in six of the seven seeds, in one of them marginally (seed 48). In seed 43 it is not. The values of one seed are printed above the panels of Fig. 11 and Fig. 13.

**Figure 13:**
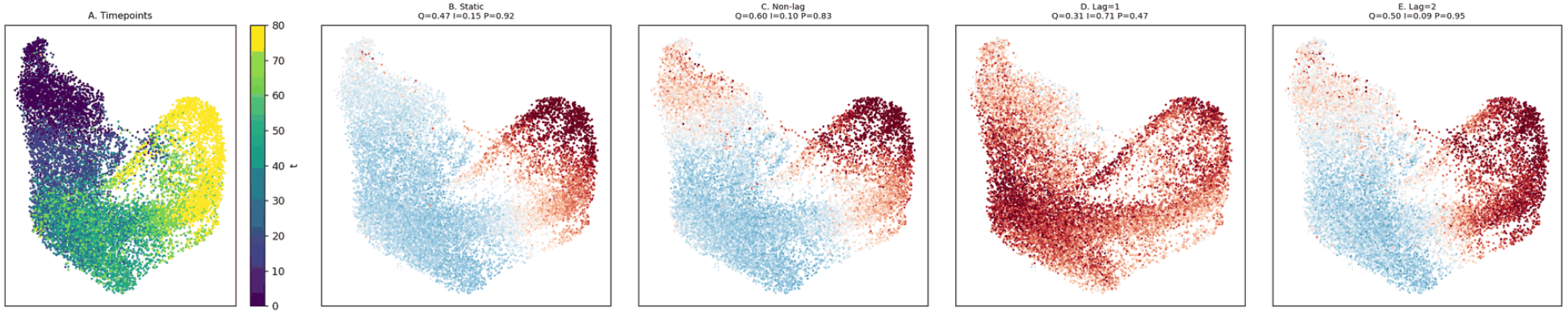
MKI67 query, reciprocal to Fig. 11. Static and non-lagged retrieval are already proliferative-dominant; lag 1 transiently loses proliferative specificity in favour of the intermediate compartment, and lag 2 restores it.

**Figure 14:**
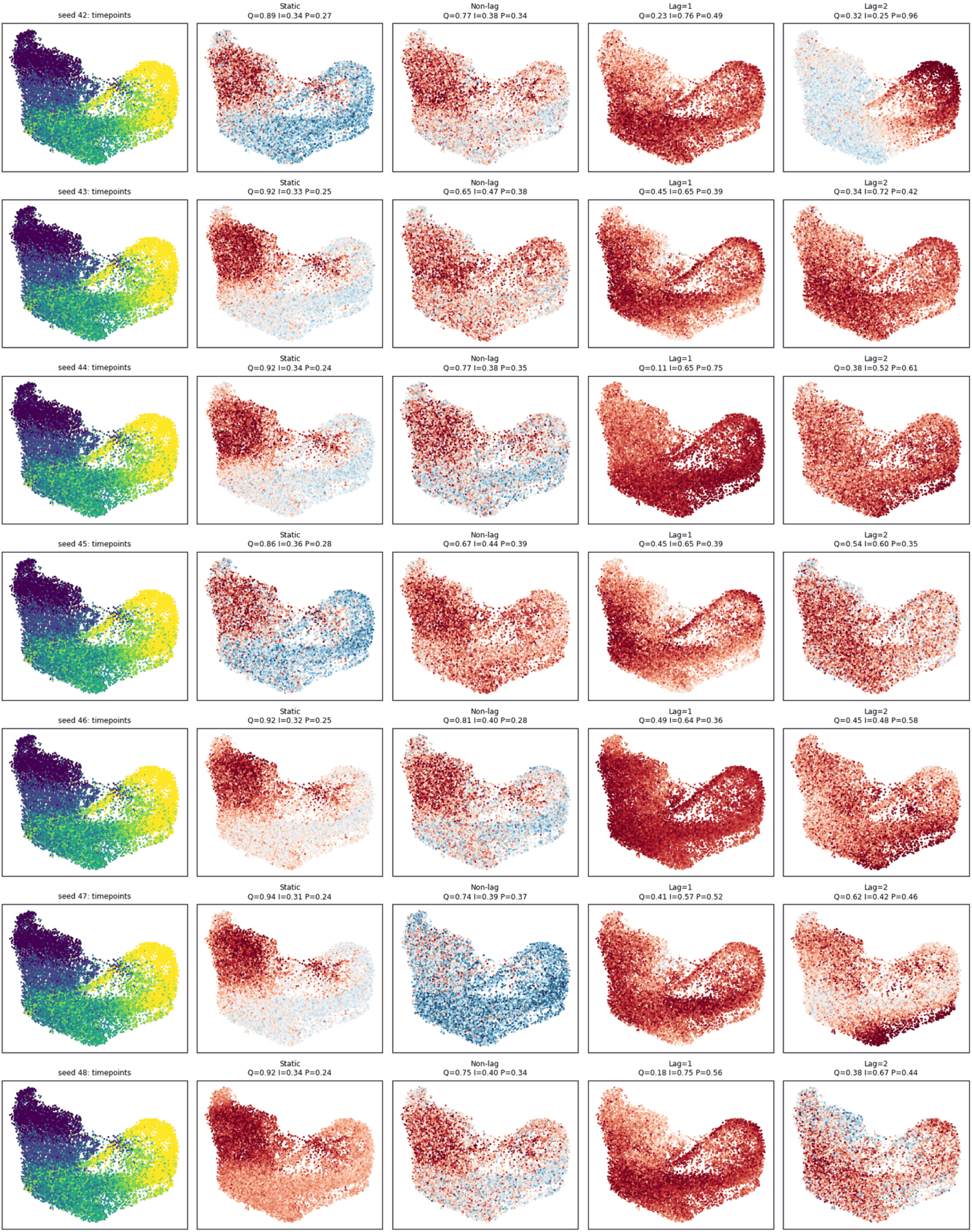
CD44 query across 7 GCN training seeds (42–48). Each row shows Timepoints, Static, Non-lag, Lag=1 and Lag=2 program scores for one seed; AUROC (Q/I/P) is reported per panel.

### MKI67

The reciprocal MKI67 query (Fig. 13) is proliferative-dominant at the first two levels, dips transiently toward intermediate at lag 1 and returns to proliferative at lag 2. Since MKI67 enters the pseudotime, it is not an independent test.

### Other query genes

The progression of HSPA1B is compared with 16 other query genes in the same pipeline and the same seven training seeds (Table 20). The set was fixed before the lagged results were computed: the ten genes with the highest AUROC of their expression for quiescent cells among the genes expressed in at least 20% of the cells (without mitochondrial and ribosomal genes), and the markers of the progenitor state of rhabdomyosarcoma reported by Danielli et al. [4] that are present in the data (MEOX2 and OGN are not expressed and POSTN is not in the data). The first step, from quiescent at the static level to intermediate at lag 1, holds in most training seeds for the genes whose static retrieval is quiescent. The second step, to proliferative at lag 2, is query-dependent: it holds in all seeds for HSPH1 and in most seeds for HSPA1B, DNAJB1 and TCEAL9, and in a minority of seeds or not at all for the others. HSPA1B, DNAJB1 and HSPH1 belong to the same heat-shock family and are therefore not independent. CD44, a progenitor marker, reaches the proliferative state at lag 2 in some seeds (Fig. 14) but passes through all three states in only two seeds. The other progenitor markers (FN1, COL1A1, EGFR, THY1) are not associated with the quiescent cells of this annotation, so they cannot serve as quiescent queries.

### Baseline on the same bins and the OT channel

Two further retrievals are scored in the same way (Table 21). The baseline describes each gene only by its mean expression in the six bins and ranks all other genes by the lagged cosine similarity of these profiles to the query gene. For HSPA1B it stays on the quiescent state at lags 0 and 1 and does not reach the proliferative state at lag 2, where the three states are scored nearly alike. The OT residual retrieves genes aligned with the quiescent state at lag 1 in six of the seven training seeds and at lag 2 in five. The proliferative state is the best-matching state at lag 1 only in seed 42 and at lag 2 in none of the seeds (proliferative AUROC at most 0.42 at lag 2; Fig. 15), so the OT channel does not trace the quiescent→intermediate→proliferative progression, which is why the KL channel is used for this analysis. The baseline is shown for seed 42, and the OT channel for seven training seeds (Fig. 15).

**Figure 15:**
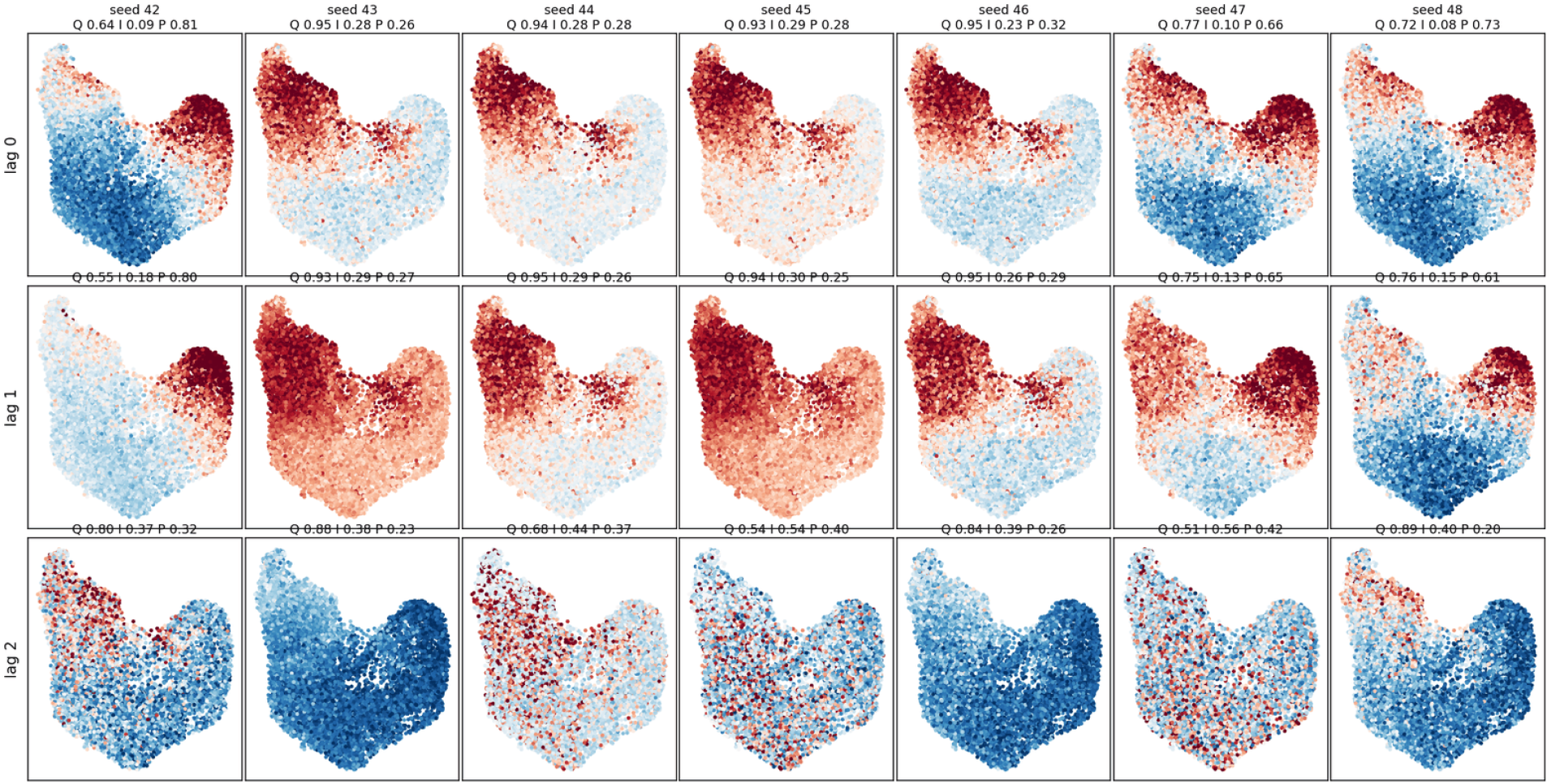
HSPA1B query, OT channel, across seven DeepSets training seeds (42–48; columns). Rows: lag 0, lag 1 and lag 2 in the forward reading. Colour: Scanpy score genes of the top 30 genes of the retrieval; AUROC against quiescent (Q), intermediate (I) and proliferative (P) cells is given above each panel.

## References

[1] Yanir Buznah, Tal Ishon, and Uri Shaham. PRISM-GEP: Viewing single-cell expression through the lens of topic modeling. Bioinformatics, 42(10):btag728, 2026.

[2] Feng Chen and Chunhe Li. Reconstructing gene network structure and dynamics from single cell data. Bioinformatics, 41(11):btaf598, 2025.

[3] Nicolas Courty, Rémi Flamary, Devis Tuia, and Alain Rakotomamonjy. Optimal transport for domain adaptation. IEEE Transactions on Pattern Analysis and Machine Intelligence, 39(9):1853–1865, 2017.

[4] Sara G. Danielli, Yun Wei, Michael A. Dyer, Elizabeth Stewart, Heather Sheppard, Marco Wachtel, Beat W. Schäfer, Anand G. Patel, and David M. Langenau. Single cell transcriptomic profiling identifies tumor-acquired and therapy-resistant cell states in pediatric rhabdomyosarcoma. Nature Communications, 15:6307, 2024.

[5] David DeTomaso and Nir Yosef. Hotspot identifies informative gene modules across dimensions of variation in single-cell data. Cell Systems, 12(5):446–456.e9, 2021.

[6] Michael B. Eisen, Paul T. Spellman, Patrick O. Brown, and David Botstein. Cluster analysis and display of genome-wide expression patterns. Proceedings of the National Academy of Sciences, 95:14863–14868, 1998.

[7] Luz Garcia-Alonso, Christian H. Holland, Mahmoud M. Ibrahim, Denes Turei, and Julio Saez-Rodriguez. Benchmark and integration of resources for the estimation of human transcription factor activities. Genome Research, 29(8):1363–1375, 2019.

[8] Ulysse Herbach. Harissa: stochastic simulation and inference of gene regulatory networks based on transcriptional bursting. arXiv preprint arXiv:2309.05112, 2023.

[9] Calley L. Hirsch, Zeynep Coban Akdemir, Li Wang, Gowtham Jayakumaran, Dan Trcka, Alexander Weiss, J. Javier Hernandez, Qun Pan, Hong Han, Xueping Xu, Zheng Xia, Andrew P. Salinger, Marenda Wilson, Frederick Vizeacoumar, Alessandro Datti, Wei Li, Austin J. Cooney, Michelle C. Barton, Benjamin J. Blencowe, Jeffrey L. Wrana, and Sharon Y. R. Dent. Myc and SAGA rewire an alternative splicing network during early somatic cell reprogramming. Genes & Development, 29(8):803–816, 2015.

[10] Sui Huang, Gabriel Eichler, Yaneer Bar-Yam, and Donald E. Ingber. Cell fates as high-dimensional attractor states of a complex gene regulatory network. Physical Review Letters, 94(12):128701, 2005.

[11] Geert-Jan Huizing, Gabriel Peyré, and Laura Cantini. Optimal transport improves cell–cell similarity inference in single-cell omics data. Bioinformatics, 38(8):2169–2177, 2022.

[12] Vân Anh Huynh-Thu, Alexandre Irrthum, Louis Wehenkel, and Pierre Geurts. Inferring regulatory networks from expression data using tree-based methods. PLoS ONE, 5(9):e12776, 2010.

[13] Masato Ishikawa, Seiichi Sugino, Yoshie Masuda, Yusuke Tarumoto, Yusuke Seto, Nobuko Taniyama, Fumi Wagai, Yuhei Yamauchi, Yasuhiro Kojima, Hisanori Kiryu, et al. RENGE infers gene regulatory networks using time-series single-cell RNA-seq data with CRISPR perturbations. Communications Biology, 6(1):1290, 2023.

[14] Thomas N. Kipf and Max Welling. Variational graph auto-encoders. In NeurIPS Workshop on Bayesian Deep Learning, 2016. arXiv:1611.07308.

[15] Thomas N. Kipf and Max Welling. Semi-supervised classification with graph convolutional networks. In International Conference on Learning Representations (ICLR), 2017.

[16] Dylan Kotliar, Adrian Veres, M. Aurel Nagy, Shervin Tabrizi, Eran Hodis, Douglas A. Melton, and Pardis C. Sabeti. Identifying gene expression programs of cell-type identity and cellular activity with single-cell RNA-Seq. eLife, 8:e43803, 2019.

[17] Evgeny Krivosheev, Mattia Atzeni, Katsiaryna Mirylenka, Paolo Scotton, and Fabio Casati. Siamese graph neural networks for data integration, 2020. arXiv:2001.06543.

[18] Solomon Kullback and Richard A. Leibler. On information and sufficiency. Annals of Mathematical Statistics, 22(1):79–86, 1951.

[19] Peter Langfelder and Steve Horvath. WGCNA: an R package for weighted correlation network analysis. BMC Bioinformatics, 9:559, 2008.

[20] Hyo-Jun Lee, Yoonji Chung, Ki Yong Chung, Yong-Kook Kim, Jea-Hyun Lee, Youn Jung Koh, and Seung Hwan Lee. Use of a graph neural network to the weighted gene co-expression network analysis of Korean native cattle. Scientific Reports, 12:9854, 2022.

[21] Patrick Lewis, Ethan Perez, Aleksandra Piktus, Fabio Petroni, Vladimir Karpukhin, Naman Goyal, Heinrich Küttler, Mike Lewis, Wen-tau Yih, Tim Rocktäschel, Sebastian Riedel, and Douwe Kiela. Retrieval-augmented generation for knowledge-intensive NLP tasks. In Advances in Neural Information Processing Systems, volume 33, pages 9459–9474, 2020.

[22] Sedigheh Mahdavi, Shima Khoshraftar, and Aijun An. Dynamic joint variational graph autoencoders. In Machine Learning and Knowledge Discovery in Databases: ECML PKDD 2019, volume 1167 of Communications in Computer and Information Science, pages 385–401. Springer, 2020.

[23] Yann Maugé and Elias Ventre. CardamomOT: a mechanistic optimal transport-based framework for gene regulatory network inference, trajectory reconstruction and generative modeling. PLOS Computational Biology, 22(10):e1014838, 2026.

[24] Thomas Moerman, Sara Aibar Santos, Carmen Bravo González-Blas, Jaak Simm, Yves Moreau, Jan Aerts, and Stein Aerts. GRNBoost2 and Arboreto: efficient and scalable inference of gene regulatory networks. Bioinformatics, 35(12):2159–2161, 2019.

[25] Mineto Ota, Jeffrey P. Spence, Tony Zeng, Emma Dann, Alexander Marson, and Jonathan K. Pritchard. Causal modelling of gene effects from regulators to programs to traits. Nature, 650:399–408, 2026.

[26] Aditya Pratapa, Amogh P. Jalihal, Jeffrey N. Law, Aditya Bharadwaj, and T. M. Murali. Benchmarking algorithms for gene regulatory network inference from single-cell transcriptomic data. Nature Methods, 17(2):147–154, 2020.

[27] Rihao Qu, Xiuyuan Cheng, Esen Sefik, Jay S. Stanley III, Boris Landa, Francesco Strino, Sarah Platt, James Garritano, Ian D. Odell, Ronald Coifman, Richard A. Flavell, Peggy Myung, and Yuval Kluger. Gene trajectory inference for single-cell data by optimal transport metrics. Nature Biotechnology, 43(2):258–268, 2025.

[28] Jolene S. Ranek, Wayne Stallaert, J. Justin Milner, Margaret Redick, Samuel C. Wolff, Adriana S. Beltran, Natalie Stanley, and Jeremy E. Purvis. Feature selection for preserving biological trajectories in single-cell data. Nature Communications, 15:2765, 2024.

[29] Noah El Rimawi-Fine, Adam Stecklov, Lucas Nelson, Mathieu Blanchette, Alexander Tong, Stephen Y Zhang, and Lazar Atanackovic. Simulation-free structure learning for stochastic dynamics. arXiv preprint arXiv:2510.16656, 2025.

[30] Clara Savary, Léa Luciana, Paul Huchedé, Arthur Tourbez, Claire Coquet, Maëlle Broustal, Alejandro Lopez Gonzalez, Clémence Deligne, Thomas Diot, Olivier Naret, et al. Fusion-negative rhabdomyosarcoma 3D organoids to predict effective drug combinations: A proof-of-concept on cell death inducers. Cell Reports Medicine, 4(12):101339, 2023.

[31] Geoffrey Schiebinger et al. Optimal-transport analysis of single-cell gene expression identifies developmental trajectories in reprogramming. Cell, 176(4):928–943, 2019.

[32] Eran Segal, Michael Shapira, Aviv Regev, Dana Pe’er, David Botstein, Daphne Koller, and Nir Friedman. Module networks: identifying regulatory modules and their condition-specific regulators from gene expression data. Nature Genetics, 34:166–176, 2003.

[33] William F. Sharpe. A simplified model for portfolio analysis. Management Science, 9(2):277–293, 1963.

[34] Itay Tirosh et al. Dissecting the multicellular ecosystem of metastatic melanoma by single-cell RNA-seq. Science, 352(6282):189–196, 2016.

[35] Tsz Pan Tong and Jun Pang. FlowGRN: Scalable and dropout-robust gene regulatory network inference via flow matching-based trajectory reconstruction. In Proceedings of the 16th ACM International Conference on Bioinformatics, Computational Biology, and Health Informatics, pages 1–8, 2025.

[36] Dénes Türei, Alberto Valdeolivas, Lejla Gul, Nicolàs Palacio-Escat, Manuel Klein, Olga Ivanova, Máté Ölbei, Attila Gábor, Fabian Theis, Dézső Módos, Tamás Korcsmáros, and Julio Saez-Rodriguez. Integrated intra- and intercellular signaling knowledge for multicellular omics analysis. Molecular Systems Biology, 17(3):e9923, 2021.

[37] Elias Ventre. Reverse engineering of a mechanistic model of gene expression using metastability and temporal dynamics. In Silico Biology, 14(3-4):89–113, 2022.

[38] Elias Ventre, Ulysse Herbach, Thibault Espinasse, Gérard Benoit, and Olivier Gandrillon. One model fits all: combining inference and simulation of gene regulatory networks. PLOS Computational Biology, 19(3):e1010962, 2023.

[39] C. H. Waddington. The Strategy of the Genes. George Allen & Unwin, 1957.

[40] Zeev Weizmann and Elias Ventre. Gene Program Explorer (RMS Explorer). Zenodo, version 1.0.0. 10.5281/zenodo.20845824, 2026. Computer software.

[41] Xiaohan Xing, Fan Yang, Hang Li, Jun Zhang, Yu Zhao, Mingxuan Gao, Junzhou Huang, and Jianhua Yao. Multi-level attention graph neural network based on co-expression gene modules for disease diagnosis and prognosis. Bioinformatics, 38(8):2178–2186, 2022.

[42] Manzil Zaheer, Satwik Kottur, Siamak Ravanbakhsh, Barnabas Poczos, Ruslan Salakhutdinov, and Alexander J. Smola. Deep Sets. In Advances in Neural Information Processing Systems, volume 30, 2017.

[43] Wenjun Zhao, Erica Larschan, Björn Sandstede, and Ritambhara Singh. Optimal transport reveals dynamic gene regulatory networks via gene velocity estimation. PLOS Computational Biology, 21(5):e1012476, 2025.

